# Generalized cell phenotyping for spatial proteomics with language-informed vision models

**DOI:** 10.1101/2024.11.02.621624

**Authors:** Xuefei (Julie) Wang, Rohit Dilip, Ahamed Raffey Iqbal, Yuval Bussi, Caitlin Brown, Elora Pradhan, Yashvardhan Jain, Kevin Yu, Shenyi Li, Martin Abt, Katy Börner, Leeat Keren, Yisong Yue, Ross Barnowski, David Van Valen

## Abstract

We present DeepCell Types, a novel approach to cell phenotyping for spatial proteomics that addresses the challenge of generalization across diverse datasets with varying marker panels collected across different platforms. Our approach utilizes a transformer with channel-wise attention to create a language-informed vision model; this model’s semantic understanding of the underlying marker panel enables it to learn from and adapt to heterogeneous datasets. Leveraging a curated, diverse dataset named Expanded TissueNet with cell type labels spanning the literature and the NIH Human BioMolecular Atlas Program (HuBMAP) consortium, our model demonstrates robust performance across various cell types, tissues, and imaging modalities. Comprehensive benchmarking shows that our method outperforms existing approaches on cell-type prediction and, from the same model, predicts marker positivity competitively with a dedicated specialist; it further matches manual expert gating and adapts to new data with modest fine-tuning, well past what baselines reach when trained from scratch. This work equips the spatial proteomics community with a single, continuously improvable phenotyping model that generalizes to new marker panels and can be fine-tuned efficiently when needed. We release both DeepCell Types and Expanded TissueNet as open-source resources.

## 1 Introduction

Understanding the structural and functional relationships within tissues is a central challenge in basic and translational research. Recent advances in multiplexed imaging have expanded the number of transcripts and proteins that can be quantified simultaneously (*1, 2, 3, 4, 5, 6, 7, 8, 9, 10*), enabling large-scale analysis of human tissue samples. Concurrently, deep learning models that integrate image and natural-language information have advanced a range of biomedical imaging applications (*11, 12, 13, 14*). However, a critical question persists: how can these innovative methods be harnessed to transform the vast amounts of data generated by multiplexed imaging into meaningful biological insights?

In this paper, we present a language-informed vision model that addresses the problem of generalized cell phenotyping in spatial proteomics data. Although modern spatial proteomics platforms generate increasingly rich datasets, significant challenges in analyzing and interpreting them at scale remain. Unlike flow cytometry or single-cell RNA sequencing, tissue imaging is performed with intact specimens. Thus, to extract single-cell data, individual cells must be identified — a task known as cell segmentation — and the resulting cells must be examined to determine their cell type and which markers they express — a task known as cell phenotyping. A general solution for cell phenotyping has proven more challenging for several reasons. First, it requires scalable, automated, and accurate cell segmentation, which has only recently become available (*15, 16, 17, 18, 19*). Second, imaging artifacts, including staining noise, marker spillover, and cellular projections, pose a formidable challenge to phenotyping algorithms (*20,21,22*). Third, general phenotyping algorithms must handle the substantial differences in marker panels, cell types, and tissue architectures across experiments. Each new dataset often has a different number of markers, each with its own distinct meaning.

Existing approaches to meet this challenge range from conventional methods that require manual gating and clustering (*23, 24, 25*) to more recent machine learning-based solutions (*26, 27, 20, 22, 21, 28*). QuPath (*25*) and histoCAT (*24*) are comprehensive interactive platforms that enable users to analyze cells based on their marker expressions and spatial characteristics, often incorporating lightweight clustering or classification tools. FlowSOM (*23*) is a commonly used clustering algorithm based on self-organizing maps. Two prominent probabilistic models, Celesta (*26*) and Astir (*27*), enable automated cell phenotyping by incorporating prior knowledge of cell type marker mappings, bypassing the need for labeled data. Celesta (*26*) employs a Markov Random Field with a mean-field approximation, while Astir (*27*) is built on variational inference with Gaussian Mixture Models. Supervised approaches like MAPS (*22*) (a feed-forward neural network), Stellar (*21*) (a graph neural network), and CellSighter (*20*) (a convolutional neural network) offer the advantage of accurate and specific cell type identification when labeled data is available. Complementing these, Nimbus (*28*) provides a deep learning model for predicting marker positivity, which can then be used with clustering algorithms like FlowSOM (*23*) for cell type identification. While these tools highlight the advances the field has made, existing methods face significant challenges in scaling for three reasons. First, many of them rely on human intervention, which places a fundamental limit on their ability to scale to big data. Second, these methods cannot handle the wide variability in marker panels that exist across experiments. Some methods require labeling and re-training for new datasets. Even when transfer learning is possible (*20, 21, 22*), the target dataset must share significant similarities in the marker panel with the source dataset. Third, most existing methods have been developed for data collected from a single imaging platform. Bridging this gap requires a versatile model that can be trained on multiple datasets and perform inference on new data with unseen markers. To this end, we developed an end-to-end cell phenotyping model that learns from and generalizes to diverse datasets, regardless of their specific marker panels. Our approach was twofold. First, we curated and integrated a large, diverse set of spatial proteomics data, sourced from the literature and the NIH HuBMAP consortium. Through a human-in-the-loop framework, experts then carried out quality control and optional annotation on fields of view (FOVs); we term the resulting labeled dataset Expanded TissueNet. Second, we developed a new deep learning method, DeepCell Types (DCT), that learns to phenotype cells across these diverse data. It combines vision and language encoders, so it can draw on both raw marker images and the semantics of the language describing markers and cell types. A transformer with channel-wise attention integrates this visual and linguistic information, removing the dependence on specific marker panels. When trained on Expanded TissueNet, our method outperforms existing approaches on cell-type prediction across diverse cell types, tissue types, and spatial proteomics platforms, and from the same model predicts marker positivity competitively with a dedicated specialist. Its panel-agnostic architecture then lets DeepCell Types generalize beyond that benchmark. It matches the neighborhood structure recovered by manual expert gating and adapts to new data with modest fine-tuning, well past what baselines reach when trained from scratch. Both Expanded TissueNet and DeepCell Types are made available through the DeepCell software library with permissive open-source licensing.

## 2 Results

Our central result is that a single language-informed vision model, trained once across our composite dataset, generalizes out of the box across the heterogeneous marker panels and imaging platforms of spatial proteomics:

- We assemble Expanded TissueNet, a corpus of ∼8.7 million expert-labeled cells spanning seven imaging platforms and standardized to a common marker and cell-type vocabulary (Fig. 1).
- We build DeepCell Types, a language-informed vision model with a channel-wise transformer, trained for cross-platform generalization and class-imbalance robustness (Fig. 2).
- DeepCell Types matches or outperforms existing methods on cell-type and marker-positivity prediction, and matches manual expert gating (Fig. 3).
- Self-supervised pre-training adds only modest in-distribution gains but reaches a higher ceiling when fine-tuning to new, label-scarce datasets.

**Figure 1:**
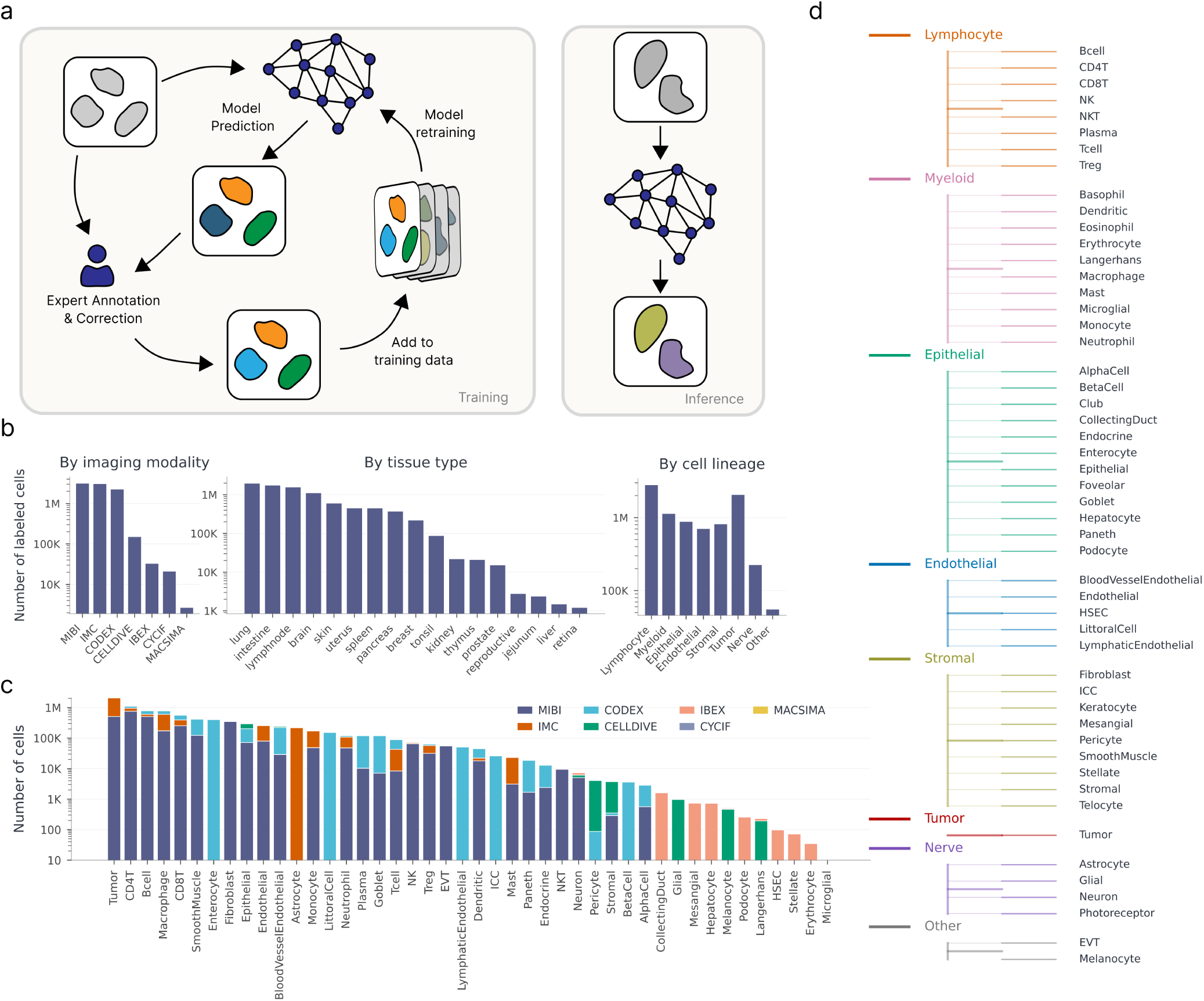
Construction of Expanded TissueNet with expert-in-the-loop labeling. **(a)** Expert-in-the-loop labeling workflow. An initial cell-type proposal — existing dataset annotation, a DeepCell Types prediction — is presented to a domain expert in DeepCell Label; the refined cell-type and marker-positivity labels are added to Expanded TissueNet, which retrains DeepCell Types and closes the loop (*Training*). At inference, the trained model instead predicts cell types directly from a raw cell patch with no expert step (*Inference*). **(b)** Composition of Expanded TissueNet by imaging modality, tissue type, and cell lineage. The labeled subset shown here spans approximately 8.7 million cells across 7 imaging platforms, 17 tissue types, and 8 lineages. **(c)** Modality composition per cell type: for each cell type (x-axis), the number of cells contributed by each imaging platform, stacked by platform. **(d)** Cell-type lineage hierarchy: 8 major lineages organized into 51 specific cell types that form the training label space; the coarser 8-lineage grouping is used for the lineage-collapsed analysis (fig. S5a).

**Figure 2:**
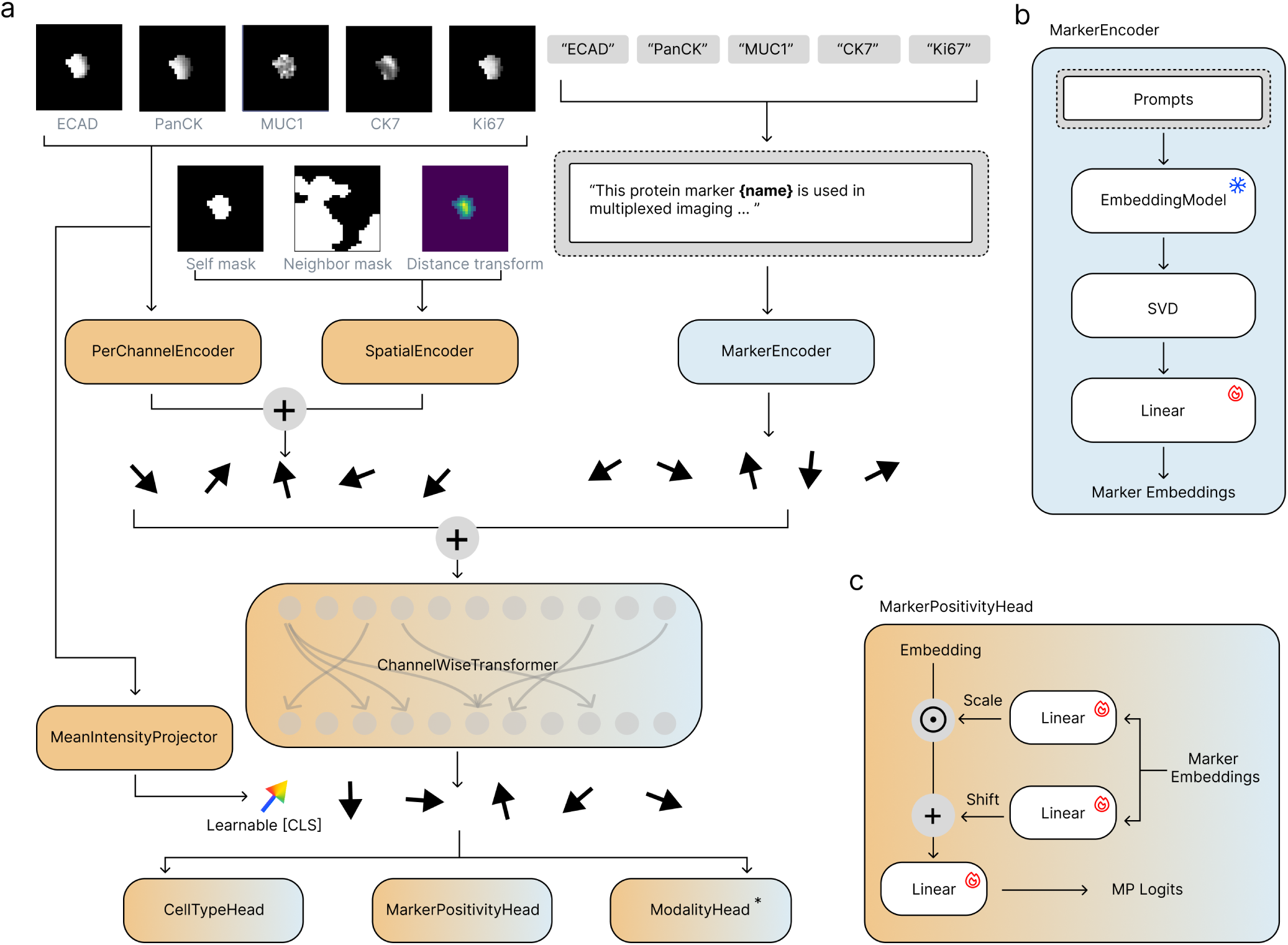
DeepCell Types: a language-informed vision model. **(a)** Overall architecture. The image lane processes each cell via a PerChannelEncoder that consumes a stack of marker patches and a parallel SpatialEncoder that consumes the self-mask, neighbor-mask, and self-mask distance transform; the two streams are combined to form one image embedding per marker channel. The language lane slots each marker name into a short biology-context template prompt (table S1) and routes it through the marker encoder to produce one text embedding per marker. Image and text embeddings are summed token-wise and consumed by a pre-norm channel-wise transformer with a learnable [CLS] token. The transformer emits joint per-marker embeddings plus the [CLS] token. A separate mean-intensity projection of the raw per-cell mean intensities is fused with the [CLS] stream and fed to a cell-type classification head, a marker-positivity head, and a modality head ModalityHead^∗^; the asterisk marks it as being trained adversarially via a gradient reversal layer (fig. S4b) so the encoder unlearns modality-specific artifacts and generalizes across imaging platforms. **(b)** Marker encoder. Each templated marker prompt is embedded by a frozen OpenAI text-embedding-3-large model into a 3072-d vector; per-marker embeddings are pre-compressed with SVD and then passed through a trainable linear layer that learns the marker embedding consumed by the rest of the model. **(c)** Marker-positivity head. The per-marker transformer outputs are conditioned by the marker embeddings through a FiLM scale-and-shift operation (one trainable scale and shift linear layer per marker), then passed through a final linear layer that produces a per-marker positivity logit. Conditioning the head per marker allows the decision boundary to be marker-adaptive.

**Figure 3:**
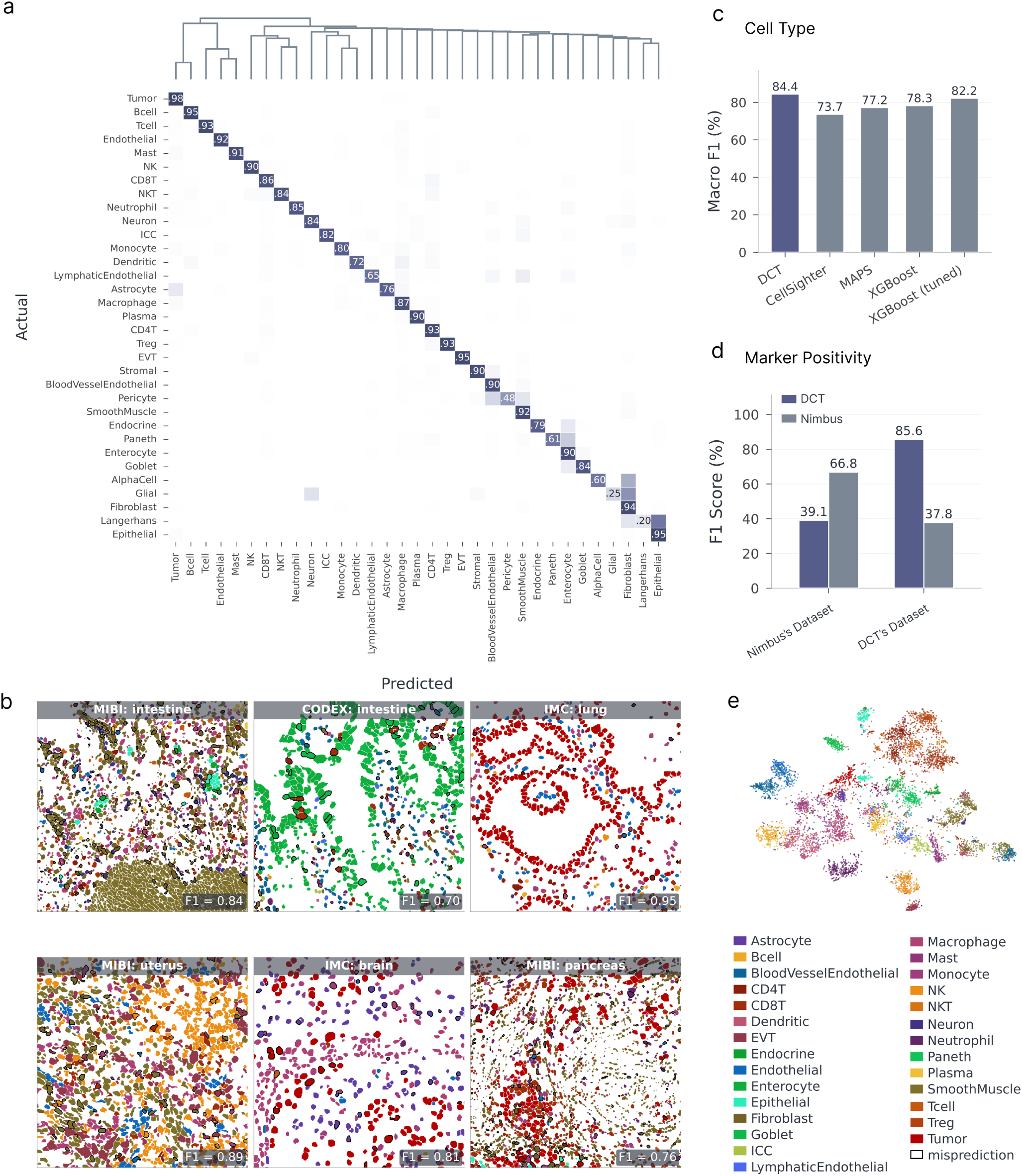
Benchmarking against existing methods on the test split. **(a)** Cell-type confusion matrix on the test split (129 FOVs / 486,705 cells). **(b)** Example fields of view spanning imaging modalities and tissues (MIBI, CODEX, and IMC across intestine, lung, uterus, brain, and pancreas), with cells colored by predicted cell type; mispredictions are outlined. **(c)** Cell-type classification performance on the unleaked test split. DeepCell Types is benchmarked against XGBoost, MAPS, and CellSighter under a common data interface, marker vocabulary, class-balancing sampler, training loop, and hierarchical evaluation protocol. **(d)** Marker positivity prediction: macro F1, comparing DeepCell Types against Nimbus on two distributions — Expanded TissueNet (DeepCell Types’ native) and the Pan-Multiplex Gold Standard (Nimbus’s native). Each model is strongest on its own distribution; per-marker breakdown and FiLM decision curves are in fig. S7. **(e)** Latent-space visualization. Two-step NCA (*56*) → t-SNE (*57*) projection (*58*) of DeepCell Types [CLS] embeddings, colored by cell type. Cell-type clusters dominate, indicating the representation is organized around biology rather than imaging platform; the same projection colored by modality and tissue is in fig. S5h–i.

### 2.1 DeepCell Label enables scalable construction of Expanded TissueNet

To capture the diversity of marker panels, cellular morphologies, tissue heterogeneity, and technical artifacts across the field, we compiled data from published sources (*29,30,31,32,33,34,35,36,37,38,39,40,41,21,42,43,44,45*) and unpublished data deposited in the HuBMAP data portal. The dataset contains 2,153 labeled FOVs and approximately 8.7 million labeled cells drawn from more than a dozen contributing sources. For each dataset, we collected raw images, corresponding channel names, and cell type labels (when available), and resized each dataset to a standard resolution of 0.5 microns per pixel (mpp). We standardized marker names and cell types across all datasets to enable cross-dataset comparison, integration, and analysis, and organized cell types by lineage (Fig. 1d).

We then performed whole-cell segmentation with Mesmer (*15*) and mapped any existing cell type labels to the resulting cell masks. When quality labels did not exist, we generated them through a human-in-the-loop labeling framework that leveraged expert labelers (Fig. 1a). To establish marker positivity labels, experts manually gated the mean signal intensity for each marker and cell type within each dataset, drawing on their knowledge of marker-cell type associations. To accelerate this step, we extended DeepCell Label (*46*), our cloud-based software for distributed image labeling, to the cell type labeling task. DeepCell Label allows users to visualize images, analyze marker intensities, refine segmentation masks, and annotate cell types.

The resulting labeled dataset, Expanded TissueNet, consists of approximately 8.7 million labeled cells, spanning seven imaging platforms: Imaging Mass Cytometry (IMC) (*3*), CO-Detection by indEXing (CODEX) (*2*), Multiplexed Ion Beam Imaging (MIBI) (*1*), Iterative Bleaching Extends Multiplexity (IBEX) (*5*), MICS (MACSima Imaging Cyclic Staining) (*4*), Multiplexed immunofluorescence with Cell DIVE^TM^ technology (Leica Microsystems, Wetzlar, Germany), and Cyclic Immunofluorescence (CycIF) (*47*), with the majority of data coming from the first three. The dataset covers 17 common tissue types and 51 specific cell types across 8 broad cell lineages (Fig. 1b,d). Beyond aggregate scale, Expanded TissueNet is diverse by construction: cell-type presence, channel co-occurrence, per-FOV cell counts, and per-FOV panel size all vary substantially across the contributing datasets (fig. S1), motivating a panel-agnostic architecture rather than a fixed-panel one. Marker discriminability is summarized in fig. S3. Across all datasets, we cataloged 278 unique protein markers, with an average of 28.5 markers per FOV.

To generate single-cell image crops for training, we extracted a 32×32 patch centered on each cell from each marker image, capturing the cell and its surrounding neighborhood. These images were augmented with a three-channel spatial-context input: a binary self-mask delineating the central cell (1’s for pixels inside the cell and 0’s for pixels outside), a binary neighbor-mask capturing surrounding cells within the patch, and a normalized distance transform of the self-mask (the Euclidean distance of each interior pixel to the cell boundary, scaled to [0, 1] by its maximum) that supplies a soft, sub-cellular shape cue peaking at the cell interior.

### 2.2 DeepCell Types is a language-informed vision model with a channel-wise transformer, trained for cross-platform generalization and class-imbalance robustness

We developed DeepCell Types, a cell phenotyping model that learns from and adapts to diverse datasets with different marker panels. Our model’s backbone consists of three main components: a visual encoder, a language encoder, and a channel-wise transformer (Fig. 2a).

#### Visual Encoder

The visual encoder turns each cell into one embedding per marker channel using two parallel Convolutional Neural Network (CNN) branches. A per-channel branch encodes the 32×32 image patch of each marker independently, while a shared spatial-context branch encodes the cell’s self-mask, neighbor-mask, and distance transform; the two are combined into a single per-channel embedding. These embeddings capture staining patterns, cell morphology, marker expression levels, and neighborhood context.

#### Language Encoder

To incorporate semantic understanding of markers and cell types, we employ a language encoder built on a large language model (LLM) that generates embedding vectors for each name (Fig. 2b). Each marker or cell type name is wrapped in a short biology-context template prompt (table S1), and the resulting prompt is passed to a frozen LLM embedder, which produces a high-dimensional embedding vector. This approach leverages knowledge about the biology of markers and cell types, enabling our model to understand the meaning behind each channel.

#### Channel-wise Transformer

To allow our model to generalize across marker panels, we use a transformer module with channel-wise attention. This module adds a marker’s image embedding with its corresponding language embedding and applies self-attention to the combined representation (Fig. 2a). This self-attention operation is applied across all the markers, similar to how a human might look across the marker images to interpret a stain. While transformers for sequence data typically include a positional encoding, we did not include one. This design choice preserved the length-and-order invariance of self-attention (*48*) and allowed our model to process inputs from diverse marker panels without modification, in contrast to fixed-channel CNNs of the CellSighter / MAPS family that hard-code a channel order and input depth (fig. S4c). We prepended a learnable [CLS] token to represent the overall cell and to aggregate information across all channels. This architecture fuses visual and linguistic information, letting our model discern cross-channel correlations together with each marker’s biological significance.

We train this architecture with five complementary design choices that jointly improve cross-platform generalization and robustness to Expanded TissueNet’s severe class imbalance.

#### Focal classification

We feed the [CLS] token embedding from the final transformer layer into a multi-class classification head, which is supervised by a focal cross-entropy loss (*49*) over the cell-type label set. The focal weighting assigns higher importance to difficult examples, which are disproportionately drawn from the long-tailed rare classes of Expanded TissueNet (fig. S2).

#### FiLM-conditioned marker positivity

Marker positivity is predicted by a marker-conditioned MLP head (Materials and Methods, marker-positivity head) that consumes the per-marker transformer outputs and is modulated channel-by-channel by the marker’s text embedding via Feature-wise Linear Modulation (FiLM) (*50*) (Fig. 2c), letting the decision boundary adapt to each marker’s expression statistics. This head is supervised by a binary cross-entropy (BCE) loss against per-cell marker-positivity labels.

#### Class-balanced sampling and a decoupled two-stage head

Expanded TissueNet is heavily class-imbalanced, so we draw training cells with a weighted sampler whose per-cell weight is proportional to the inverse square root of the cell’s class frequency (Materials and Methods, class-balanced sampling). This sampler is necessary for the channel-wise backbone to learn good features for rare classes, but leaving it on for the cell-type head over-fires those same rare classes. We therefore use a decoupled two-stage recipe for all reported cell-type results: the sampler-trained backbone is frozen, and a residual-MLP cell-type head is retrained on the natural (sampler-off) class distribution.

#### Mean-intensity residual

To complement the image-and-text representation with an explicit quantitative cue, we add a *mean-intensity [CLS] residual*. For each cell, we compute the mean raw intensity of every present marker inside the self-mask, scatter the values into a fixed-length vector indexed by the model’s global marker vocabulary, project that vector through a small zero-initialized MLP, and add the result to the post-transformer [CLS] embedding before the cell-type head (Materials and Methods, mean-intensity residual). Because the per-channel CNN is FOV-normalized and InstanceNorm-pooled, the image branch is intentionally invariant to per-marker absolute intensity, which nevertheless remains the dominant feature used by traditional intensity-gating workflows. Routing the per-cell mean intensities through a vocabulary-indexed residual recovers this cue without disturbing the panel-agnostic transformer path.

#### Adversarial modality-invariance

During the development of Expanded TissueNet, we noted significant class imbalances with respect to imaging modalities (Fig. 1c). To prevent the model from overfitting to any single modality, we implemented an auxiliary task that encourages platform-invariant representations. To do so, we added a classification head that takes [CLS] token embeddings and predicts imaging modalities, coupled with a gradient reversal layer (*51*). During backpropagation, the gradients into the encoder are reversed, so the encoder is pushed to unlearn modality-specific differences (fig. S4b). The resulting model focuses on the underlying biological characteristics of cells rather than platform-specific artifacts (Fig. 3e, fig. S5h).

### 2.3 DeepCell Types matches or outperforms existing methods and manual expert gating

#### Cell-type prediction

We benchmarked DeepCell Types against existing methods across imaging modalities, tissue types, and cell types (Fig. 3c). Fig. 3b shows example fields of view spanning these inputs, and the full cell-type confusion matrix on the test split is shown in Fig. 3a. To test whether the learned representation captures cross-platform biology rather than platform-specific artifacts, we visualize DeepCell Types’ [CLS] embeddings. The resulting projection (Fig. 3e) shows that cells cluster by cell type rather than by imaging modality (fig. S5h), indicating that the representation is organized around biology rather than acquisition platform.

To assess head-to-head performance against existing methods on the test split, we benchmarked DeepCell Types against three baselines: XGBoost (*52*), a tree-boosting algorithm; MAPS (*22*), an MLP-based approach over per-cell mean marker intensities; and CellSighter (*20*), a CNN-based approach over multi-channel cell patches. To ensure that comparisons reflect modeling choices rather than implementation differences, we re-implemented every baseline against a common data interface, marker vocabulary, train/validation/test split, and evaluation protocol on Expanded TissueNet; full implementation details and code paths are reported in the Baselines subsection of Materials and Methods. In this evaluation, differences in marker panels are flagged as missing data for XGBoost and zero-padded for MAPS and CellSighter. As shown in Fig. 3c, our language-informed vision model reaches 84.4% macro cell-type F1 on the held-out test split, outperforming XGBoost (78.3%), MAPS (77.2%), and CellSighter (73.7%). This XGBoost baseline uses the package’s default hyperparameters; an additional Optuna hyperparameter sweep pushes this strong baseline further to 82.2%, yet the tuned model still trails by 2.2 points. This shows that integrating language and vision information handles the heterogeneous marker panels of spatial proteomics data more effectively than approaches that ignore marker semantics. Because Expanded TissueNet is heavily class-imbalanced, we report macro F1 as our primary metric. Per-dataset macro F1 on the bottom 40 of 129 test FOVs (fig. S6), the lineage-collapsed confusion matrix (fig. S5a), per-class precision–recall (fig. S5b), and macro F1 stratified by imaging modality, lineage, and tissue (fig. S5c–e), together with model calibration and prediction-confidence diagnostics (fig. S5f,g), are reported in the supplement.

#### Marker-positivity prediction

We also benchmarked our model’s marker-positivity prediction against Nimbus on both Expanded TissueNet and the Pan-Multiplex Gold Standard, a dataset curated by Nimbus (*28*) (Fig. 3d). Each model performs best on its native distribution: on Expanded TissueNet our model reaches 85.6% macro F1, versus 37.8% for Nimbus. On the Pan-Multiplex Gold Standard, Nimbus reaches 66.8% macro F1 against our 39.1%. This asymmetry tracks training distribution, not a general ranking: Expanded TissueNet spans DeepCell Types’ native diversity of platforms and panels, while the Pan-Multiplex Gold Standard was assembled by the Nimbus authors and matches Nimbus’s own training data. Each model is the stronger choice on the distribution it was built for, rather than one uniformly outperforming the other. The per-marker F1 breakdown, the learned per-marker threshold distribution, and marker-specific decision curves for representative markers (CD45, CD206, SMA) are in fig. S7.

#### Matching manual expert gating

Using raw image data as the primary input is a substantive shift from traditional cell phenotyping, which relies on per-cell mean intensity values extracted from segmented cells followed by overclustering and expert rule-based mapping. That pipeline requires a domain expert’s time for every new dataset and struggles with the artifacts of spatial proteomics data. On a shared FOV, DeepCell Types reaches 0.93 macro F1 against ground truth end-to-end, versus 0.71 for the mean-intensity-overcluster-then-expert-merge pipeline (fig. S8b). The two workflows also organize tissue similarly at the neighborhood level. Comparing *k* = 15 cellular-neighborhood maps (*53*), they reach 78.6% cell-wise label agreement and a low mean Jensen–Shannon divergence of 0.128 bits between their neighborhood-composition distributions (fig. S8c); the rule-based map is less accurate cell-for-cell yet broadly congruent with DeepCell Types in local neighborhood structure. DeepCell Types produces these predictions without any per-dataset expert gating effort, saving on the order of hours of hands-on expert time. Inference itself is inexpensive: per-channel prediction time scales linearly with the number of cells, at 0.04 ± 0.01 ms per cell per channel (fig. S9d).

### 2.4 Self-supervised pre-training yields modest gains but a higher fine-tuning ceiling

Given the size and diversity of Expanded TissueNet and the recent successes of self-supervised pre-training in vision and language, a natural hypothesis was that an initial self-supervised stage would yield a stronger representation than supervised training alone. We tested this directly by implementing a masked-marker pre-training objective: for each cell, we randomly mask a fraction of valid marker channels at the input and train the model to predict each masked channel’s mean per-cell intensity from the remaining channels and the corresponding language embeddings (Materials and Methods, masked-marker pre-training). After pre-training, we fine-tune the full model end-to-end with the standard cell-type and marker-positivity losses.

We first confirmed that the pretext task itself is learnable: scored on held-out cells, the pre-training backbone recovers masked marker positivity well above an untrained baseline (fig. S9a), so the pre-training objective does capture genuine cross-marker structure. Yet relative to training the supervised objective from scratch, that pre-training yields only a modest downstream gain on both cell-type classification and marker-positivity prediction (fig. S9b).

The pre-trained backbone’s practical payoff instead surfaces in the label-scarce regime, when adapting to a single target dataset with limited labels. On the Keren MIBI triple-negative breast cancer dataset (*33*), we fine-tune this masked-marker-pretrained backbone end-to-end on labeled subsets of *N* = 8 to 533 FOVs and compare against XGBoost and CellSighter trained from scratch on the same subsets (fig. S9c). In the very-low-label regime (*N* ≤ 16), a simple, more regularized model like XGBoost gives a moderate lift from scratch, while fine-tuning a large pretrained model from only a handful of examples is not yet worth its cost (e.g. 71.2% vs 47.3% at *N* = 8). But once enough labels are available to amortize that cost, DeepCell Types reaches a higher ceiling than either baseline attains on the same labels: it overtakes XGBoost by *N* = 32 and, with the full labeled set, reaches 90.2% macro F1 — 8.9 points above XGBoost (81.3%) and 17.6 points above CellSighter (72.6%).

## 3 Discussion

This work addresses a critical challenge in spatial proteomics: the need for automated, generalizable, and scalable cell phenotyping tools. Our approach directly tackled the fundamental issue of variability in marker panels across different experiments, enabling learning from multiple data sources. Trained on a diverse dataset, our model phenotypes cells accurately across the modalities, tissues, and marker panels present in Expanded TissueNet, matching or outperforming existing methods and manual expert gating. For marker positivity, each model is strongest on its native distribution: DeepCell Types leads on Expanded TissueNet while Nimbus leads on its own Pan-Multiplex Gold Standard, reflecting complementary specializations rather than a uniform ranking.

DeepCell Types departs from traditional cell phenotyping approaches, which originate from non-spatial single-cell technologies like scRNA-Seq and flow cytometry and rely on mean intensity values extracted from segmented cells, often failing to cope with the artifacts of spatial proteomics data — a contrast we quantify directly in the Results. DeepCell Types nevertheless fuses per-cell mean intensities as an auxiliary signal through the mean-intensity [CLS] residual (Materials and Methods, mean-intensity residual), recovering the quantitative gating cue without depending on it as the sole input. Our work adds to the growing body of evidence that image-centric approaches are more robust to segmentation errors, noise, and signal spillover (*20, 28*).

A key innovation in our model is its use of language to improve generalization. By incorporating textual information about markers, the model draws on biological knowledge that extends beyond the training data. This pairing of visual and linguistic information lets DeepCell Types integrate evidence across a broad set of experimental data and outperform traditional machine-learning approaches. The architecture can also train on multiple datasets simultaneously, exploiting marker correlations across datasets — a capability existing methods lack. It can therefore improve continuously as new data emerges, consolidating the field’s knowledge into a single, increasingly comprehensive model. Consistent with this view, adding a self-supervised pre-training stage on top of the supervised objective yielded only modest further gains (Results), suggesting that the diversity of labeled data in Expanded TissueNet, rather than the training objective, is already the dominant driver of the model’s cross-marker representations.

Despite these advances, our method has limitations that merit discussion. First, like any data-driven system the model’s performance is optimal within the domain of training data and is expected to degrade on samples too far out-of-domain. Expanded TissueNet does not exhaust all imaging modalities and tissue types. Cell type and imaging modality are also coupled: for cell types contributed by a single platform, the adversarial modality-invariance objective (Materials and Methods, gradient-reversal head) cannot fully separate biology from platform-specific artifacts. Hence, even though we showed promising improvement compared to existing methods, datasets that substantially diverge from Expanded TissueNet may require additional labeling and fine-tuning to achieve adequate performance, though fine-tuning still reaches a higher ceiling than training a baseline from scratch once enough labels are available to amortize its cost (Results). To help users assess and correct such misalignment, we release an automated preprocessing-adaptation procedure as the preproc-adapt agent skill (Wang et al. (*54*)), which corrects resolution, intensity-scaling, and marker-naming mismatches without touching model weights (Supplementary Methods). We anticipate our labeling software and human-in-the-loop approaches to labeling will enable these efforts and facilitate the collection and labeling of increasingly diverse data, further improving model performance. As demonstrated here and previously (*15, 16*), integrated data and model development is viable for consortium-scale deep learning in the life sciences. Second, DeepCell Types is limited to the cell types in its training ontology (table S2), which does not capture the wide variety of additional, tissue-specific cell types across human biology; extending coverage to this broader diversity is an important direction for future work. Last, our model is trained on cell patches and cannot access the full image context. While this constraint on the effective receptive field can promote generalization, it may also limit accuracy. Features at longer length scales like functional tissue units and anatomical structures can provide contextual information to further aid cell type determination. Beyond these limitations, the same panel-agnostic architecture that handles the heterogeneous marker panels of spatial proteomics here is a natural fit for image-based spatial transcriptomics or marker-aware cell segmentation, and extending it to those settings is a promising next step.

In summary, DeepCell Types gives the community a panel-agnostic, scalable framework for cell phenotyping that generalizes across imaging platforms and marker panels. Released alongside Expanded TissueNet as open, continuously improvable resources, the two turn cell phenotyping into a shared, cumulative effort for the field, rather than a problem every lab must re-solve alone for every new panel.

## 4 Materials and Methods

### 4.1 Data preprocessing and standardization

We implemented several preprocessing steps that were applied to each marker image independently to ensure consistent and high-quality input for our model. To standardize image resolution, all datasets were resampled to 0.5 mpp; raw marker intensities were resampled with bilinear interpolation, while integer segmentation masks were resampled with nearest-neighbor interpolation and without anti-aliasing. The resampling factor is stored as a per-FOV attribute in the dataset: of the 2,772 preprocessed FOVs, 53.1% were downsized to reach 0.5 mpp, 46.8% were upsized, and the remaining 0.1% were already at native 0.5 mpp. We thresholded the images using the 99.9th percentile of all non-zero values per channel per FOV and then applied min-max normalization. Processed images were saved in zarr format, enabling chunked, compressed storage and fast loading. We also standardized marker names and cell type labels across datasets, establishing a canonical name for each marker and cell type, identifying and mapping all alternative names accordingly (e.g., “Pan-Keratin” and “PanCK” to a single canonical marker name; “Treg” and “regulatory T cell” to a single canonical cell type).

Cell segmentation was performed using the Mesmer algorithm (*15*). We identified nuclear channels (e.g., DAPI, Histone H3) and cytoplasm/membrane channels (e.g., Pan-Keratin, CD45) to generate accurate whole-cell masks. For each segmented cell, we extracted a 32×32 pixel patch centered on the cell. For cells near image boundaries, we added padding to ensure complete 32×32 patches. To capture segmentation information, we constructed a per-cell spatial-context tensor by stacking the self-mask, the neighbor-mask, and the distance transform of the self-mask, yielding a tensor of shape (3, 32, 32). The per-channel raw images were stacked into a tensor of shape (*C,* 1, 32, 32), where *C* is the number of marker images varying by dataset. We padded the channel dimension to *C_max_* = 80 with the sentinel value −1 (chosen to lie outside the [0, 1] normalized intensity range so padded slots are unambiguously distinguishable from real signal) to allow processing by regular neural networks. A binary padding mask of (*C_max_,*) is also generated and propagated to the transformer’s attention so that padded slots cannot be attended to. Marker positivity labels were generated by expert labelers, who manually gated the mean signal intensity for each marker and cell type within each dataset, drawing on their knowledge of marker-cell type associations.

### 4.2 Model architecture

Our model employs a hybrid architecture, combining Convolutional Neural Network (CNN) and Transformer components to effectively process both image and textual data. The backbone comprises three components — a visual encoder, a language encoder, and a channel-wise transformer — on top of which we add the task heads and auxiliary branches described below. These modules, together with the training procedure (Model training), realize the five design choices highlighted in the Results: focal cell-type classification, FiLM-conditioned marker positivity, class-balanced sampling with a decoupled two-stage head, the mean-intensity [CLS] residual, and adversarial modality-invariance.

#### Visual encoder

The visual encoder is composed of two parallel CNN branches that produce a single per-channel embedding. (i) A *spatial-context encoder* consumes the (3, 32, 32) stack of self-mask, neighbor-mask, and self-mask distance transform with three stride-2 2D convolution layers (3→32→64→64 channels, kernel size 3, padding 1), each followed by *batch normalization* and SiLU activation, and yields a 64-dimensional spatial embedding shared across all channels. We deliberately use BatchNorm rather than InstanceNorm in this branch because per-sample instance normalization combined with the global pool would zero out the per-cell mean of the spatial-context maps and discard cell-size and cell-density information that we want the encoder to preserve. (ii) A *per-channel encoder* processes each marker channel independently with a stride-2 stem convolution (kernel 3, padding 1), three residual blocks (each two stride-1 convolutions with kernel 3, padding 1), global average pooling, and a final linear projection from the base-channel width to 128 dimensions, all using *instance normalization* and SiLU, producing a 128-dimensional per-channel embedding. InstanceNorm is required in this branch because zero-padded channels (used to align panels of varying width to *C_max_*) would corrupt BatchNorm running statistics; per-sample normalization makes each channel’s representation independent of how many real channels accompany it in the batch. We reshape the per-channel input from (*B, C_max_,* 1, 32, 32) to (*B* · *C_max_,* 1, 32, 32), allowing a single CNN to process all channels regardless of their marker correspondence. This design ensures the model remains agnostic to specific marker representations. The 64-dimensional spatial embedding is broadcast across channels and concatenated with the 128-dimensional per-channel embedding, then linearly projected to produce image embeddings of size (*B, C_max_,* 256). The per-channel ResNet uses a base channel width of 48.

#### Language encoder

For the language encoder, we used OpenAI’s text-embedding-3-large model (*55*) as the LLM embedder. Each marker name is wrapped in a short biology-context template prompt (“*The protein biomarker* ⟨*name*⟩ *is used in multiplexed immunofluorescence imaging for cell type identification in tissue samples.*”) and each cell-type name is wrapped in an analogous prompt (“*The cell type* ⟨*name*⟩ *as identified in multiplexed immunofluorescence imaging of human tissue samples.*”); the templates are listed in table S1. The marker and cell-type prompts are batched into a single call to the embedder, which produces a 3072-dimensional embedding vector per prompt. The embeddings are reduced via truncated singular value decomposition (SVD) fit jointly and then linearly mapped to 256 dimensions to match the image embeddings, and finally *L*_2_-normalized so that each marker token enters the transformer on the unit sphere. The *L*_2_ normalization is applied at both training and inference time (no train/eval mismatch) and serves two purposes: it stabilizes the scale of the language signal added to the image embedding regardless of dataset-specific variance in the SVD basis, and it makes the marker-conditioned FiLM head’s ***γ****_c_,* ***β****_c_* predictions a function of a marker’s *direction* in semantic space rather than its magnitude.

#### Channel-wise transformer

To merge information contained in the image and text embeddings, we fed them into a transformer module that uses channel-wise attention. A learnable [CLS] token was prepended to the embedding tensor to represent the entire cell. The transformer module comprises 4 pre-norm encoder layers with a hidden dimension of 256, 8 attention heads, and a feed-forward dimension of 1024 (dropout *p* = 0.1 applied to attention outputs and feed-forward activations). We employ a padding mask to exclude dummy channels, ensuring the model focuses only on real channels (fig. S4a). This flexible mechanism accommodates varying numbers of input channels.

#### Cell-type classification head

For cell type classification, we extract the [CLS] token embedding and feed it into the cell-type classification head: a residual-MLP with an input projection to width 512, four residual blocks (linear, batch normalization, SiLU, dropout 0.2), and a linear readout.

#### Marker-positivity head

For marker positivity, we use a marker-conditioned head that operates on the per-channel transformer outputs **h***_c_* ∈ R*^d^*^model^. For each channel *c* corresponding to marker *m_c_* with text embedding **e***_m_*, the head computes a Feature-wise Linear Modulation (*50*): ***γ****_c_* = *σ*(*W_γ_***e***_mc_*) and ***β****_c_* = *W_β_***e***_mc_*, applies 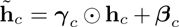, and feeds 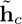 through a two-layer MLP (*d*_model_ → *d*_model_*/*2 → 1) with SiLU activation to yield a per-channel positivity logit. The scale weight *W_γ_* is initialized to zero and the scale bias to 4.0 so that *σ*(·) ≈ 0.98 at initialization, recovering an approximately marker-agnostic head at the start of training; the modulation magnitude grows as the head learns marker-specific decision boundaries. This head is supervised by a binary cross-entropy loss against per-cell marker positivity labels.

At inference time, the per-channel positivity logits are converted to binary calls with *per-marker* thresholds learned post-hoc on the training split rather than a single global 0.5 cutoff. A global threshold is miscalibrated for the long tail of low-prevalence markers, whose positivity distributions are shifted toward zero by the BCE objective, and consequently understates marker-positivity macro F1. The per-marker thresholds are obtained by sweeping each marker’s threshold on the training predictions and selecting the value that maximizes that marker’s F1. The FiLM head and the per-marker threshold sweep are two halves of the same calibration mechanism: the FiLM modulation (***γ****_c_,* ***β****_c_*) gives each marker its own decision *shape*, conditioned on the marker’s text embedding, and the post-hoc threshold sweep selects the best operating *point* on each marker-specific shape. The spread of the learned thresholds (range [0.01, 0.79], median ≈ 0.40 for the released artifact, against a default of 0.5) is direct evidence that the FiLM head produces marker-specific decision functions rather than a marker-agnostic 0.5-thresholded sigmoid, with the post-hoc sweep closing the residual operating-point gap on each marker.

#### Mean-intensity [CLS] residual

In parallel with the image-and-text path through the transformer, we provide the classification head with an explicit per-marker mean-intensity signal extracted directly from the segmented cell. For each channel *c* present in the cell, we compute the mean raw intensity inside the self-mask, 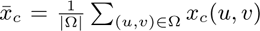, where Ω is the set of pixels inside the cell footprint. The per-cell, per-channel values {*x̅_c_*} are scattered into a fixed-length vector **m** ∈ R*^M^* indexed by the model’s global marker vocabulary (*M* = 278 for this model), with absent markers held at zero. This sparse vector is passed through a small two-layer MLP (Linear(*M* → *d*_model_) → GELU → Dropout → Linear(*d*_model_ → *d*_model_), denoted *f*_MI_) whose final layer is zero-initialized, and the result is added as a residual to the post-transformer [CLS] embedding before the cell-type head: **z̃**_CLS_ = **z**_CLS_ + *f*_MI_(**m**). The zero-initialization ensures that at the start of training the model is exactly equivalent to one without this branch, so the residual can only contribute information that improves the supervised objective. The motivation is twofold: (i) the per-channel CNN is intentionally agnostic to absolute intensity (each channel is min-max normalized per FOV and processed independently with InstanceNorm), so the model otherwise has no direct access to per-marker intensity magnitudes which remain a strong cue for traditional intensity-based phenotyping; and (ii) anchoring the marker-intensity signal to a fixed, vocabulary-indexed slot decouples the intensity cue from *C_max_* padding and channel order, mirroring how a human analyst reads quantitative gating thresholds while still interpreting the underlying images.

#### Adversarial modality-invariance head (gradient reversal)

We additionally route the [CLS] embedding through a 3-layer MLP (each hidden layer followed by LayerNorm, SiLU, and dropout; LayerNorm is used in place of BatchNorm so that activations remain comparable across small modality-imbalanced batches) that predicts the imaging modality, with a gradient-reversal layer (*51*) between the embedding and the head so that the modality loss *maximizes* the encoder’s modality confusion while the cell-type and marker-positivity losses are minimized. The intent is to discourage the encoder from latching onto platform-specific intensity statistics that do not transfer across imaging modalities, complementing the FOV-level intensity standardization performed during preprocessing.

### 4.3 Model training

#### Training objective

Our training objective is a weighted sum of three losses, one per prediction head. Cell-type classification uses a focal cross-entropy loss (*49*) with *γ* = 2.0, which up-weights difficult, often rare-class examples and improves performance on more challenging categories (the class-frequency imbalance itself is corrected by the weighted sampler, below). Marker-positivity prediction uses a binary cross-entropy loss. The reverse imaging-modality prediction (the gradient-reversal head in Model architecture) uses a cross-entropy loss with label smoothing 0.01. The cell-type and marker-positivity terms are held at weight 1.0; the modality term enters with a constant weight of 0.1 in our default configuration (set to zero to disable the head). The gradient-reversal coefficient *λ*is fixed to 1 throughout training.

#### Optimization

We train the model for 50 epochs using the AdamW optimizer with a peak learning rate of 1 × 10^−3^ and weight decay 0.01, scheduled with a OneCycleLR cosine policy with 5% warmup; the default batch size is 256 cells. Training runs under Automatic Mixed Precision on GPU by default, and gradients are clipped to a global norm of 1.0 before the optimizer step. We employ random horizontal and vertical flipping as image-level data augmentation. To encourage the model to learn robust features that are less dependent on specific markers, we randomly drop up to 8 marker channels during training (capped at 30% of valid channels per sample).

#### Class-balanced sampling

Class imbalance is addressed at the sampler level: we draw cells with a weighted random sampler whose per-cell weight is proportional to the inverse square root of the cell’s class frequency in the training split, with the per-class count floored at 1,000 before the square root (so a class with *<* 1,000 cells is treated as if it had exactly 1,000). This floor caps the amplification of single-FOV rare classes that would otherwise receive runaway per-sample weights (*>* 10^2^) and corrupt the representations of common classes. Because the sampler already corrects the imbalance, we disable per-class weights inside the focal loss when this sampler is active to avoid double-counting the correction.

#### Decoupled two-stage recipe

The headline cell-type results (Fig. 3c) use this recipe in two stages. Stage 1 is the supervised backbone training described above, with the weighted sampler on so that rare classes receive enough exposure for the channel-wise backbone to learn good features. The same sampler, however, hurts the cell-type head: it over-fires rare classes. In stage 2 we therefore freeze the entire backbone and retrain only the cell-type head on the *natural* class distribution (sampler off). Concretely, we extract the frozen backbone’s CLS embedding for every training and validation cell, standardize it to zero mean and unit variance, and retrain the head with plain (unweighted) cross-entropy, using AdamW (learning rate 1.5 × 10^−3^, weight decay 10^−4^, OneCycleLR schedule) for 50 epochs over the cached embeddings. At deployment the input standardization is folded into the head’s first linear layer, so the released model consumes the raw CLS embedding directly.

### 4.4 Masked-marker self-supervised pre-training

For the masked-marker pre-training objective, we mask 30% of valid channels per cell and use as the regression target the mean intensity of each masked channel over the cell’s segmented footprint (taken as the union of non-zero channels’ self-mask pixels, matching the per-cell mean computed inside the model). We pre-train for 20 epochs using AdamW (peak learning rate 3 × 10^−4^, batch size 256, OneCycleLR cosine schedule, 500,000 samples per epoch) on the same training data used by the supervised model. We additionally apply the standard marker-positivity BCE auxiliary loss (weight 0.5) during pre-training so that the FiLM-conditioned marker-positivity (MP) head is warm-started rather than randomly initialized at the start of fine-tuning.

To evaluate the pretext task on its own terms (fig. S9a), we score the pre-training backbone’s masked-channel intensity predictions on held-out cells against two references: an untrained backbone with randomly initialized weights (the floor), and the same backbone given every channel visible, i.e. without masking (the fully-visible reference, an upper bound since the target channel’s own intensity is then part of the input).

### 4.5 Train / validation / test split

Throughout this work we evaluate on a fixed, file-level FOV split. The split is constructed in two stages with a fixed random seed (default 42). *Stage 1* stratifies all 2,153 labeled FOVs by (imaging modality, tissue) bucket and holds out FOVs proportionally within each bucket at a 0.8/0.2 ratio, yielding 1,722 training FOVs and 431 held-out FOVs. *Stage 2* sub-partitions the 431 held-out FOVs by a 0.7/0.3 global shuffle (same seed) into a 302-FOV **validation set** used for best-epoch selection and early stopping, and a 129-FOV **test set** (486,705 cells) that is never touched by any model’s selection process. All reported test-set numbers come from this 129-FOV partition. Two corner cases are handled deterministically at Stage 1. (i) *Sole-source forcing* : any FOV that is the only field of view in the dataset containing a given cell type is forced into the training split, so that no rare-but-present class is silently dropped at training time. (ii) *Single-FOV strata*: any (modality, tissue) bucket containing exactly one FOV is also forced into training, since a held-out FOV from such a bucket would be unevaluable.

### 4.6 Evaluation protocol and treatment of rare cell types

We report cell-type performance primarily as *macro F1*, which averages the per-class F1 over cell types and weights every cell type equally regardless of its frequency. Expanded TissueNet contains a long tail of rare cell types. While we retain all 51 fine-grained cell types in our cell-type ontology (table S2) and our human-in-the-loop labeling pipeline so that they can be predicted at inference time, a subset of these labels is too rare in our current dataset to support meaningful held-out evaluation. To prevent these classes from inflating or destabilizing reported metrics, we apply a minimum-support floor of 50 at evaluation.

### 4.7 Hierarchical cell-type scoring

Several of the cell types in our ontology have well-defined parent–child relationships that the cell-type label set does not enforce explicitly. In particular, Tcell is treated as the parent of CD4T, CD8T, Treg, and NKT, and Stromal is treated as the parent of Fibroblast and Pericyte. A naïve flat (non-hierarchical) scoring penalizes a model equally for predicting CD4T when the ground-truth label is Tcell (a finer-grained refinement of the same lineage) and for predicting CD4T when the ground-truth label is Epithelial (a different lineage entirely), even though the former is a biologically reasonable, more-specific call rather than an error. To avoid this overpenalty for the Tcell and Stromal parents, we apply a *hierarchical collapsing* when scoring, alongside the flat (uncollapsed) scoring. Concretely, for cells whose ground-truth label is Tcell, predictions of any of CD4T, CD8T, Treg, or NKT are also counted as correct; for cells whose ground-truth label is Stromal, predictions of Fibroblast or Pericyte are also counted as correct. All other ground-truth labels are scored as in the flat case.

### 4.8 Latent-space visualization

To visualize the learned [CLS] embeddings (Fig. 3e, fig. S5h,i), we apply a two-step dimensionality reduction on a class-balanced 10,000-cell subsample of the test split: Neighborhood Components Analysis (NCA) (*56*) to a target dimension of 8 (100 L-BFGS iterations), followed by t-SNE (*57*) (perplexity 150, early exaggeration 4.0). This two-step scheme exposes latent-space structure without overfitting (*58*).

### 4.9 Baselines

Reimplementation was necessary because neither MAPS’s nor CellSighter’s reference implementation supports Expanded TissueNet’s heterogeneous, multi-panel marker vocabulary. The resulting harness — a single CLI shared across all four baselines — also lets future methods be benchmarked on Expanded TissueNet under the same data, splits, and metrics.

To provide a fair, reproducible point of comparison, we re-implemented each cell-type baseline — XGBoost, MAPS, and CellSighter — against the same data interface, marker vocabulary, train/validation/test split, and evaluation protocol used for DeepCell Types. All three read from the same zarr-format Expanded TissueNet dataset, consume the same per-cell feature representation (multi-channel cell-centered image patches for the image-based baseline; per-cell mean intensities aligned to the universal 278-marker vocabulary for non-image baselines, with absent markers flagged as missing, NaN, for XGBoost and zero-padded for MAPS and CellSighter), and are evaluated with the same hierarchical cell-type scoring. This makes the comparison a measure of modeling choices rather than of incidental differences in preprocessing, marker harmonization, or scoring. Where we deviate from a baseline’s reference implementation, we do so to keep the inputs and the training loop comparable; each deviation is recorded below.

We implemented XGBoost using the Python XGBoost package (https://github.com/dmlc/xgboost), with ab-sent markers explicitly indicated as missing through the data matrix interface. We report two XGBoost configurations trained on the same data, features, and split. The default baseline is fully trained on the complete training split using the package’s default hyperparameters (100 boosting rounds, at max depth 6), with early stopping on the held-out val-idation split (early stopping rounds = max(10, n estimators*/*10)). To push for an even stronger baseline, the tuned model additionally selects its hyperparameters by an Optuna (*59*) sweep over n estimators (100–1500), max depth (3–12), learning rate (0.005–0.3, log-uniform), min child weight (1–10), subsample (0.5–1.0), and colsample bytree (0.5–1.0), with early stopping via the package’s native early stopping rounds criterion on the held-out validation split, keeping the test set held out. Fig. 3c reports both the default XGBoost baseline and the hyperparameter-tuned XGBoost; the Optuna trial progression for the sweep is shown in fig. S9e.

For MAPS, we reproduced the model from the reference implementation at https://github.com/mahmoodlab/MAPS, re-implementing it end-to-end inside our training framework. We retained the reference input schema — per-cell mean intensities concatenated with the per-cell size scalar, a 279-dimensional vector — the reference MLP architecture (four 512-wide hidden layers with dropout 0.25), and the reference default optimizer (Adam, constant learning rate 10^−3^, no scheduler). We adapt three aspects to keep the inputs and model selection comparable across baselines. First, feature normalization is ((*x* − *µ*)*/σ*)*/*255: the 1*/*255 scaling is retained from the reference mahmoodlab/MAPS pipeline, and the leading per-feature *z*-score is a DCT-side adaptation that places our [0, 1]-normalized marker means and the raw-pixel cell-size scalar on a common scale before the 1*/*255 step. Second, we train for up to 500 epochs following the canonical mahmoodlab/MAPS schedule (minimum 250 epochs; early-stopping patience of 100 epochs on inner-validation loss), keeping the lowest-inner-val-loss checkpoint. Third, we select that checkpoint on a FOV-grouped 10% inner-validation split (GroupShuffleSplit) carved from the training FOVs rather than on the reported test set, so test signal never drives model selection; consequently MAPS trains on ≈ 90% of the training cells.

In addition to XGBoost and MAPS, we re-implemented CellSighter (*20*), a convolutional baseline that classifies cell types directly from multi-channel image patches. Following the reference implementation at https://github.com/KerenLab/CellSighter, we used a torchvision ResNet-50 backbone with the original ImageNet stem (a 7 × 7 stride-2 convolution followed by max-pooling) on 60 × 60 cell crops, trained from random initialization. The network input is the marker channels aligned to the universal vocabulary together with a cell (self) mask and a neighbor mask (NUM MARKERS + 2 channels). CellSighter was trained for 50 epochs with Adam at a constant learning rate of 10^−3^ and no scheduler — matching the original CellSighter recipe — with the best epoch selected by macro F1 on a FOV-grouped 10% inner-validation split carved from the training FOVs rather than on the reported test set, so that test signal never drives model selection. Like other image-based baselines, CellSighter does not natively reason about marker semantics across panels with different channel sets, so its zero-shot evaluation uses the same marker-channel zero-padding scheme noted above.

To benchmark our model’s performance in marker positivity prediction, we separately compared it to Nimbus (*28*), a pretrained deep learning model that predicts marker positivity directly, without requiring retraining. We used the released model weights and followed the reference inference recipe (https://github.com/angelolab/Nimbus-Inference), reading each FOV’s per-marker channels directly from the Expanded TissueNet zarr store and computing the Nimbus-style per-channel normalization dictionary on the fly so the inputs presented to Nimbus match the statistics it expects from its native pipeline. For evaluation on the Pan-Multiplex Gold Standard (Nimbus’s native distribution), DeepCell Types is scored under a 2-fold cross-validated, per-marker thresholding protocol so that the threshold applied to each cell is never tuned on that cell. Concretely, after matching DeepCell Types’ per-cell marker-positivity scores to the gold-standard binary activity labels on (FOV, cell, marker), we partition the gold FOVs into two folds by a fixed shuffle (seed 42) and an even split, so that the partition is at the FOV level and no cell is shared across folds. Per-marker F1 is then computed on the combined cross-validated calls and macro-averaged over the 60 shared markers using the Nimbus evaluation routine, so that DeepCell Types and Nimbus are scored by the same implementation.

## Acknowledgments

We thank Noah Greenwald, Michael Angelo, Sean Bendall, Georgia Gkioxari, Edward Pao, Uriah Israel, Ellen Emerson, and the other members of the Van Valen lab for helpful feedback and interesting discussions. We thank John Hickey and Jean Fan for contributing novel datasets.

## Author Contributions

XW and DVV conceived the project; XW, YY, and DVV developed the DeepCell Types model architecture; XW, RD, and EP developed data engineering infrastructure; XW, EP, RD, and RB curated data; YB, YJ, KB, LK contributed novel datasets and provided insight on image labeling; YB, CB, and MA performed image labeling; XW, AI, and RB performed benchmarking and validation; KY, SL, and RB developed cell phenotyping capabilities for DeepCell Label; RB oversaw software engineering and deployment within HuBMAP; XW, DVV wrote the paper; XW, DVV, LK reviewed and revised the paper; DVV supervised the project.

## Data and Materials Availability

All data needed to evaluate the conclusions in the paper are present in the paper and/or the Supplementary Materials. Source code for model inference, training and baseline reimplementation is available at https://github.com/vanvalenlab/deepcell-types. Instructions for downloading the comprehensive dataset compiled for this study, along with the pretrained model weights, are available at https://vanvalenlab.github.io/deepcell-types. The code to reproduce the figures included in this paper is available at https://github.com/vanvalenlab/DeepCellTypes-2026_Wang_et_al.

## Supplementary Materials

**Supplementary Table S1:**
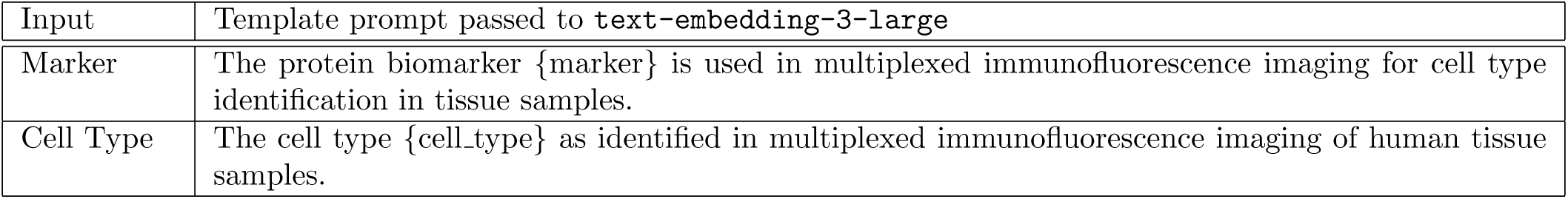
Language Encoder template prompts. Each marker or cell-type name (the bracketed slot) is substituted into the corresponding short biology-context string, and the resulting prompt is embedded directly with OpenAI text-embedding-3-large (no intermediate LLM-explainer stage). The two prompt templates are the only text-side inputs to the released embeddings; the slot-fill text comes from the marker / cell-type vocabulary used by the model.

**Supplementary Table S2:** Mapping of our cell types to HuBMAP categories based on Cell Ontology (CL).

| Our Category | Ontology ID | Ontology Name |
| --- | --- | --- |
| Tcell | CL:0000084 | T cell |
| Treg | CL:0000815 | regulatory T cell |
| CD4T | CL:0000624 | CD4-positive, alpha-beta T cell |
| CD8T | CL:0000625 | CD8-positive, alpha-beta T cell |
| NKT | CL:0000814 | mature NK T cell |
| Bcell | CL:0000236 | B cell |
| Plasma | CL:0000786 | plasma cell |
| NK | CL:0000623 | natural killer cell |
| Dendritic | CL:0000451 | dendritic cell |
| Mast | CL:0000097 | mast cell |
| Neutrophil | CL:0000775 | neutrophil |
| Basophil | CL:0000767 | basophil |
| Eosinophil | CL:0000771 | eosinophil |
| Macrophage | CL:0000235 | macrophage |
| Microglial | CL:0000129 | microglial cell |
| Langerhans | CL:0000453 | Langerhans cell |
| Monocyte | CL:0000576 | monocyte |
| Erythrocyte | CL:0000232 | erythrocyte |
| Epithelial | CL:0000066 | epithelial cell |
| CollectingDuct | CL:1001225 | kidney collecting duct cell |
| Club | CL:0000158 | club cell |
| Foveolar | CL:0002179 | foveolar cell of stomach |
| Goblet | CL:0000160 | goblet cell |
| Paneth | CL:0000510 | paneth cell |
| Enterocyte | CL:0000584 | enterocyte |
| Hepatocyte | CL:0000182 | hepatocyte |
| Keratocyte | CL:0002363 | keratocyte |
| Podocyte | CL:0000653 | podocyte |
| Endocrine | CL:0000163 | endocrine cell |
| AlphaCell | CL:0000171 | pancreatic A cell |
| BetaCell | CL:0000169 | type B pancreatic cell |
| Endothelial | CL:0000115 | endothelial cell |
| HSEC | CL:1000398 | endothelial cell of hepatic sinusoid |
| LymphaticEndothelial | CL:0002138 | endothelial cell of lymphatic vessel |
| BloodVesselEndothelial | CL:0000071 | blood vessel endothelial cell |
| LittoralCell | CL:1000397 | endothelial cell of venous sinus of red pulp of spleen |
| Stromal | CL:0000499 | stromal cell |
| Fibroblast | CL:0000057 | fibroblast |
| Stellate | CL:0000632 | hepatic stellate cell |
| SmoothMuscle | CL:0000192 | smooth muscle cell |
| Pericyte | CL:0000669 | pericyte |
| Mesangial | CL:0000650 | mesangial cell |
| ICC | CL:0002088 | interstitial cell of Cajal |
| Telocyte | NA | telocyte (no canonical CL ID) |
| Neuron | CL:0000540 | neuron |
| Glial | CL:0000125 | glial cell |
| Astrocyte | CL:0000127 | astrocyte |
| Photoreceptor | CL:0000210 | photoreceptor cell |
| Tumor | NA | NA |
| EVT | CL:0008036 | extravillous trophoblast |
| Melanocyte | CL:0000148 | melanocyte |

**Supplementary Figure S1:**
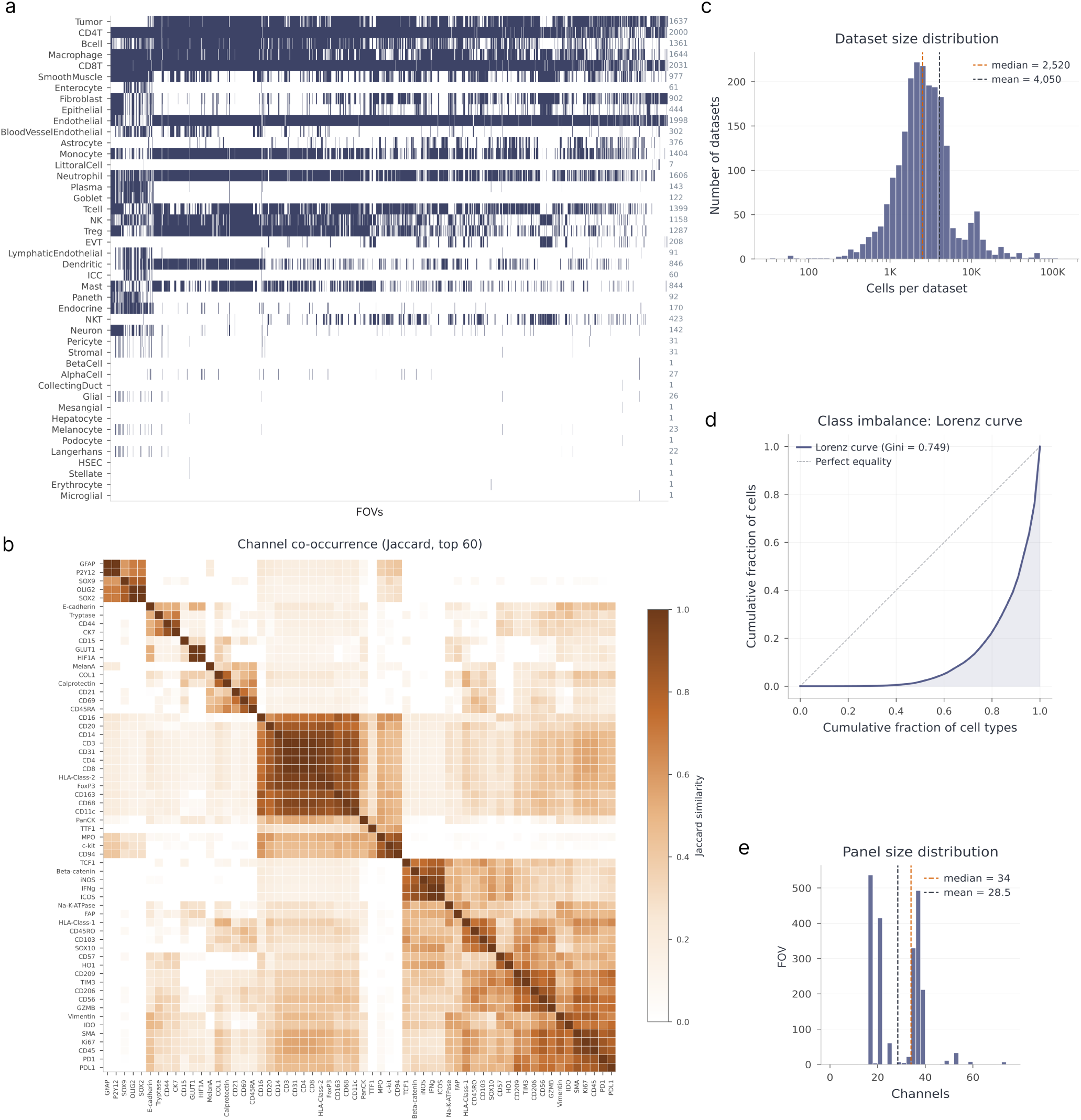
Expanded TissueNet coverage. **(a)** Cell-type presence per FOV; dark cells indicate that a given cell type is present in a given FOV. **(b)** Channel co-occurrence (Jaccard similarity) across datasets, showing which markers tend to be co-acquired in the same panel. **(c)** Per-FOV cell-count distribution across all curated FOVs. **(d)** Class imbalance curve across cell types in the dataset. **(e)** Panel-size distribution: histogram of the number of standardized markers per FOV, summarizing the panel-width heterogeneity that motivates the panel-agnostic design.

**Supplementary Figure S2:**
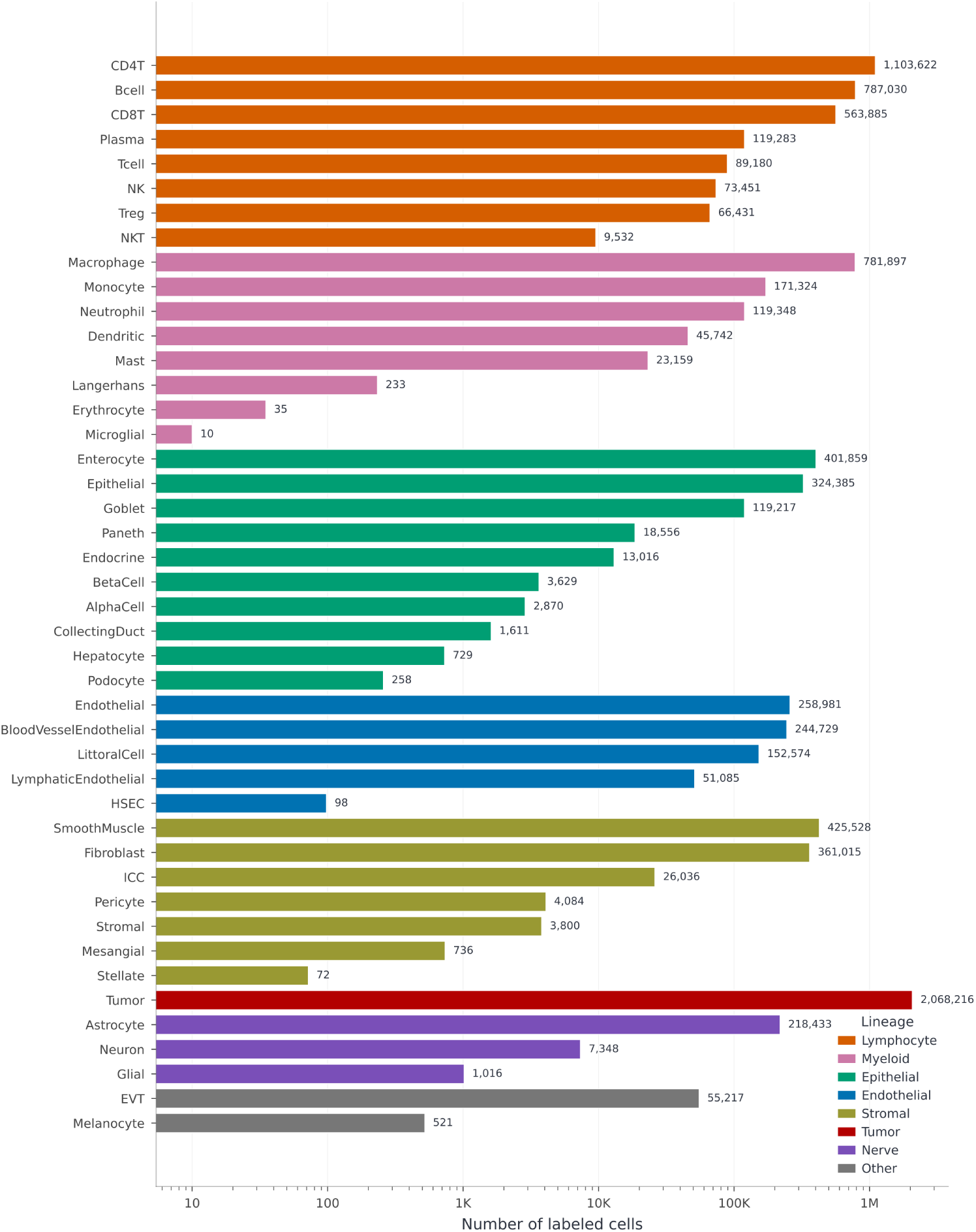
Cell-type abundance. Global cell-type frequency across the dataset (log-scaled, colored by lineage), illustrating the long-tailed distribution that motivates the focal-loss + weighted-sampler training strategy.

**Supplementary Figure S3:**
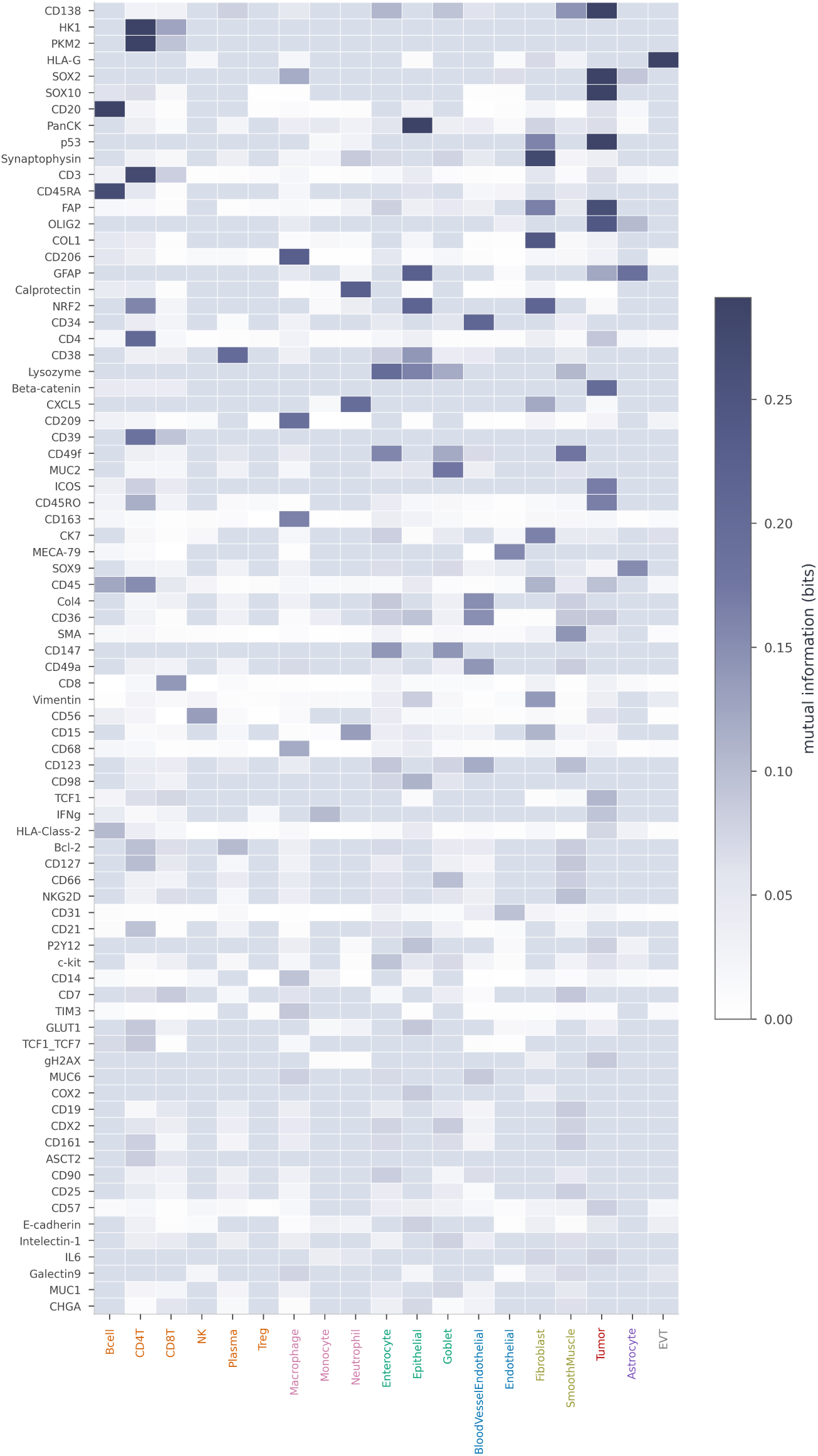
Single-channel marker discriminability. Per-cell mutual information (in bits) between each marker’s intensity and the one-vs-rest cell-type indicator. Intensities are quantile-binned (16 bins shared across cell types per row), and each entry is the plug-in mutual information *I*(*X*; *Y*) between the binned intensity *X* and the binary membership label *Y* = :ll[cell = ct]; 0 bits means intensity is uninformative about membership in that cell type. Rows (markers) are ranked by max mutual information across cell types; columns (cell types) are grouped by lineage.

**Supplementary Figure S4:**
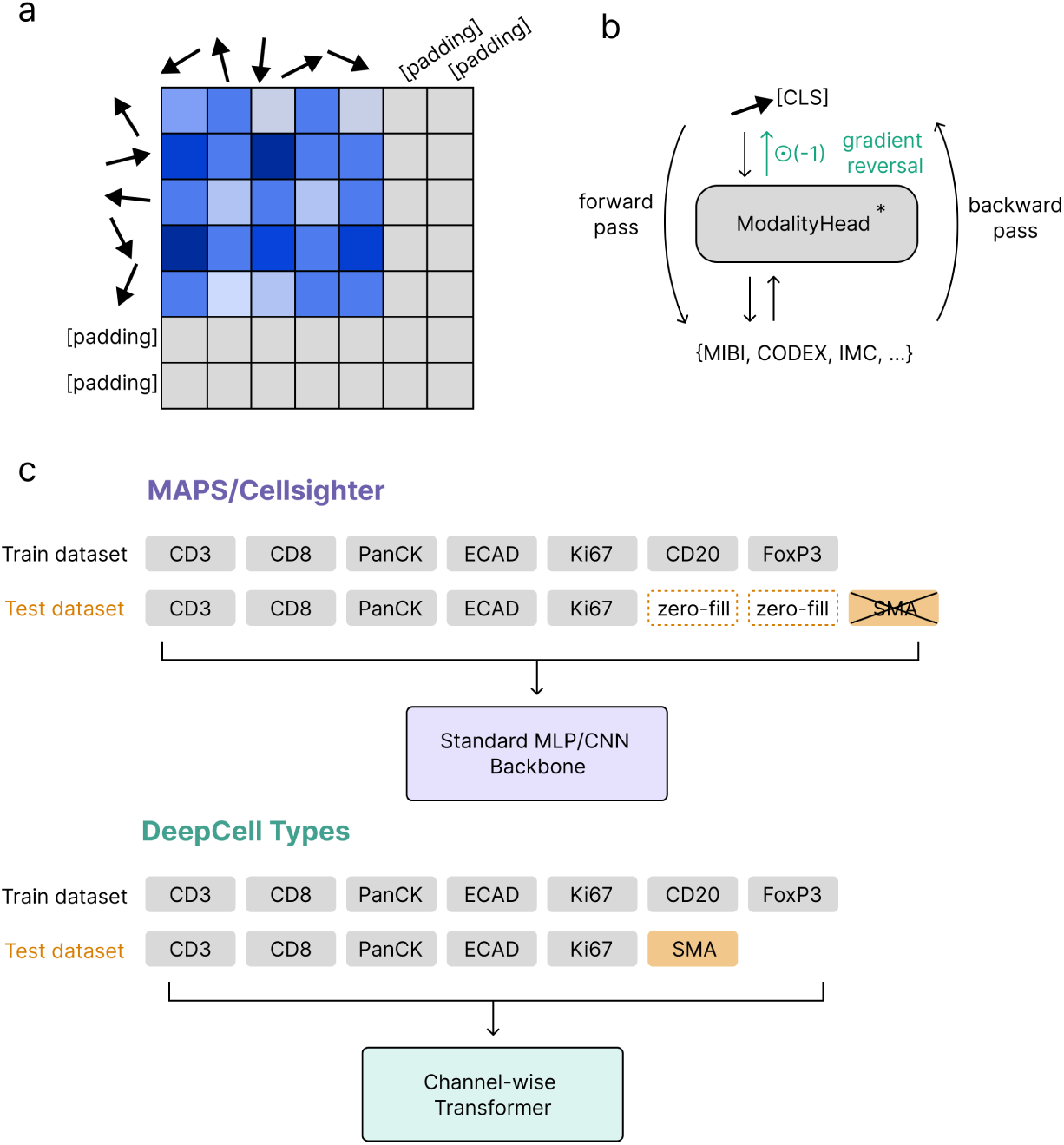
Additional architecture detail. **(a)** Channel-wise transformer attention mask. A binary padding mask is applied during self-attention so that attention is computed only over the real marker tokens in each FOV’s panel; padding slots used to align panels of varying width to *C_max_* channels are ignored. **(b)** Gradient-reversal domain-adversarial modality head. The [CLS] embedding is routed through a gradient-reversal layer and a 3-layer MLP that predicts the imaging modality. During the forward pass the head computes a modality cross-entropy loss; during the backward pass the gradient-reversal layer flips the sign of the gradient flowing into the encoder, so the encoder is pushed to *unlearn* modality-specific features. **(c)** Architectural positioning of DeepCell Types relative to a fixed-channel CNN of the CellSighter / MAPS family. Fixed-channel CNNs ingest a stacked multichannel cell patch with a hard-coded channel order and a fixed input depth, so adapting them to a new panel either requires retraining or implicit zero-padding of missing channels. DeepCell Types instead processes each marker channel independently with a shared per-channel encoder, then attends across channels in a channel-wise transformer conditioned on language-derived marker tokens; the model is therefore agnostic to panel size and marker order at both training and inference time.

**Supplementary Figure S5:**
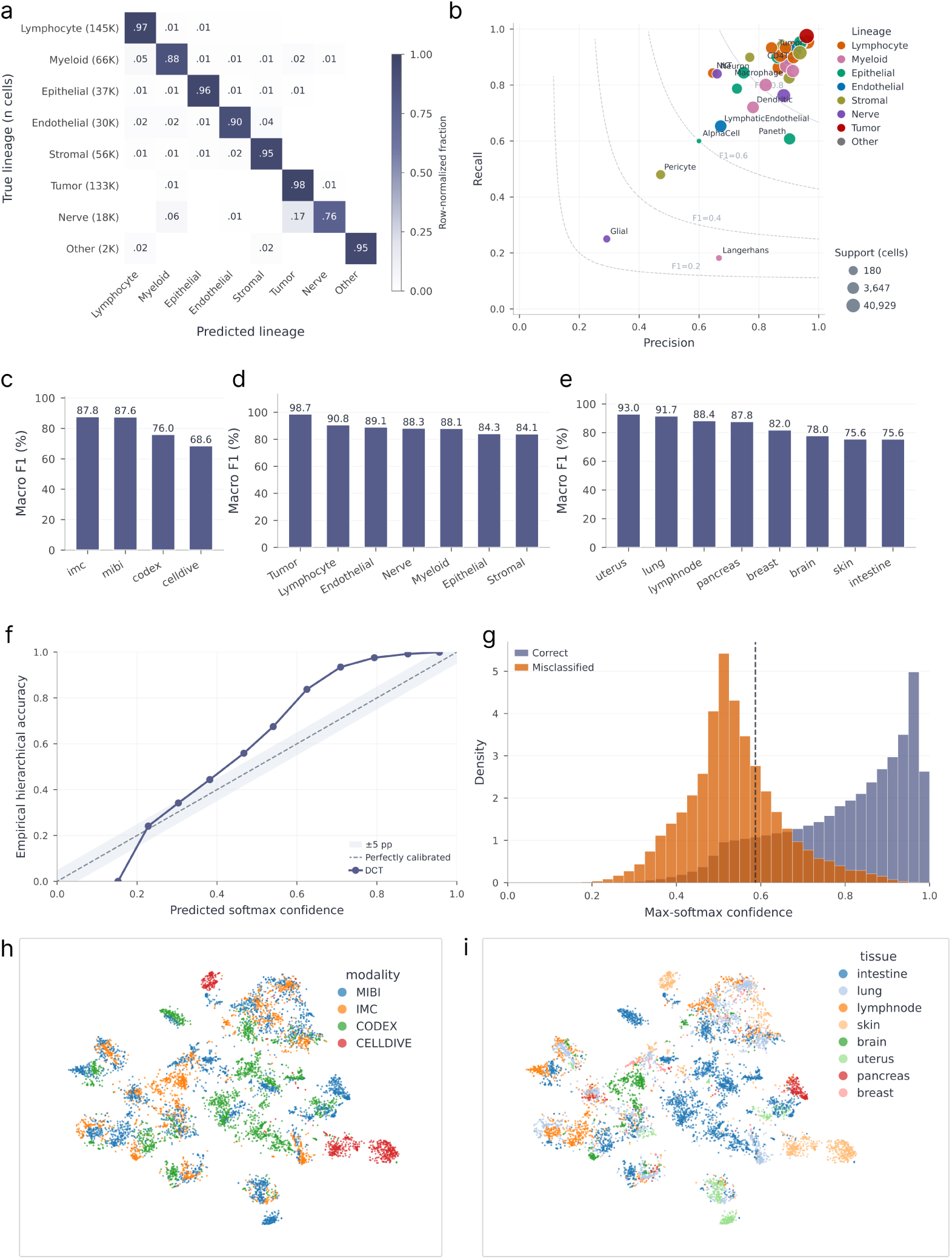
Classification diagnostics. **(a)** Lineage-collapsed confusion matrix on the test split (DeepCell Types, row-normalized) — the 8×8 coarse-grained companion to the 33×33 fine-grained panel. **(b)** Per-class precision vs. recall on the held-out test split; dot color encodes lineage, dot size encodes log(support), dashed gray curves are F1 isocontours. **(c)** DeepCell Types macro F1 on the test split, grouped by imaging modality. **(d)** DeepCell Types macro F1 grouped by true-label lineage. **(e)** DeepCell Types macro F1 grouped by tissue / organ. **(f)** Reliability (calibration) curve: empirical accuracy within confidence bins versus the model’s predicted softmax confidence, compared against the perfectly-calibrated diagonal (±5 percentage-point band shaded). Points on the diagonal are perfectly calibrated; the curve largely tracks the diagonal within the shaded band, indicating that the model’s predicted confidence is relatively well calibrated with its empirical accuracy rather than overly confident. Accuracy also rises monotonically with confidence (panel g), so the max-softmax score can be used to flag or abstain on low-confidence predictions. **(g)** Distribution of max-softmax confidence for correctly classified versus misclassified test cells; correct predictions concentrate at high confidence while misclassified cells cluster at lower, more uncertain confidence. **(h)** NCA → t-SNE projection of DeepCell Types [CLS] embeddings colored by imaging modality. Intermixed colors indicate the representation is not dominated by platform-specific features (the cell-type-colored view, Fig. 3e, organizes by biology instead; the tissue-colored view is panel i). The model uses our default configuration, with the gradient-reversal modality head active at weight 0.1 (fig. S4b). **(i)** NCA → t-SNE projection of the same embeddings colored by tissue. As with the modality view (panel h), tissues are intermixed rather than block-separated, indicating the cell-type-organized structure in Fig. 3e is not a tissue-of-origin artifact.

**Supplementary Figure S6:**
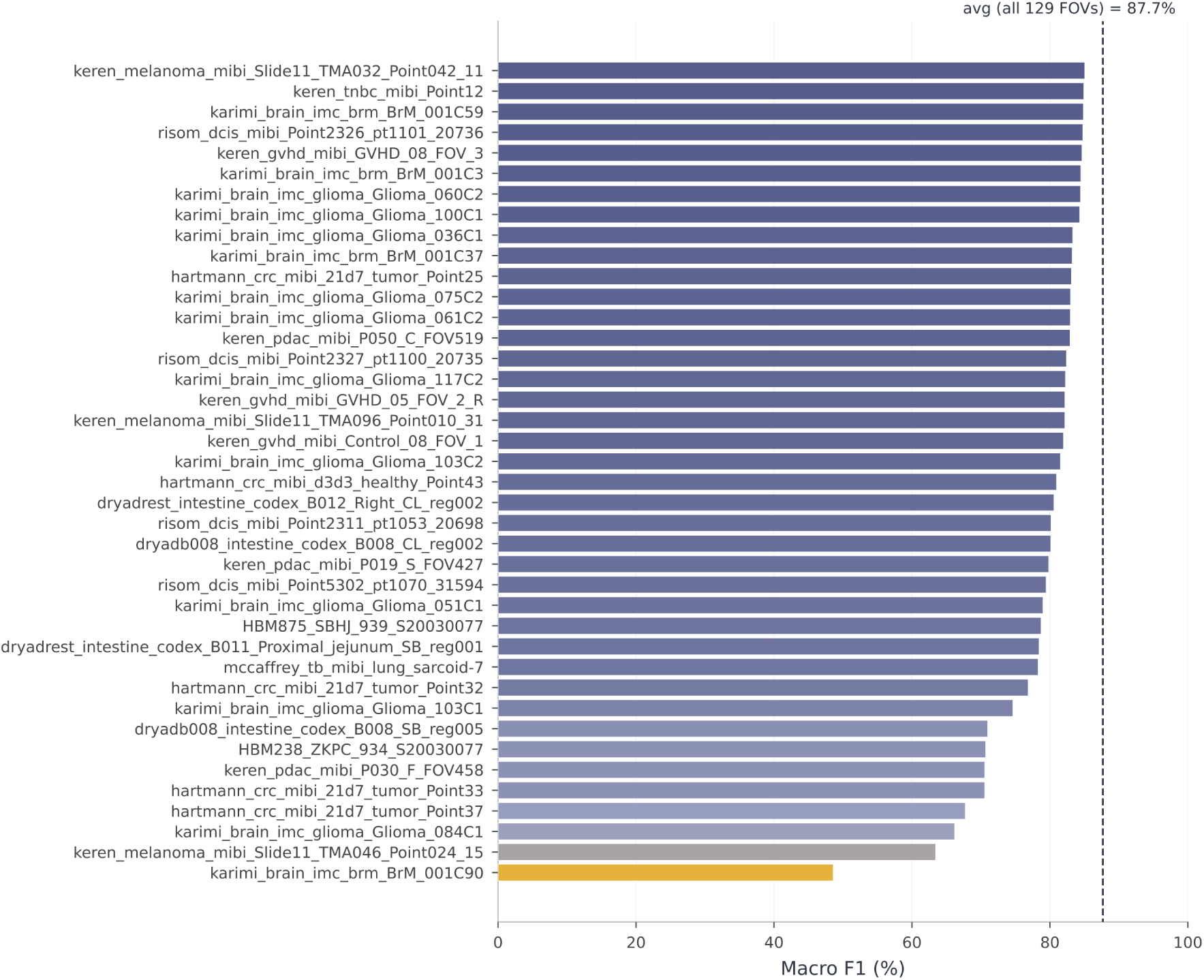
Per-dataset cell-type macro F1. Per-dataset (per-FOV) cell-type macro F1 on the test split, bottom 40 of 129 FOVs, ordered ascending. The worst-served FOVs are typically those with small panels or unusual marker sets, consistent with the panel-agnostic generalization envelope.

**Supplementary Figure S7:**
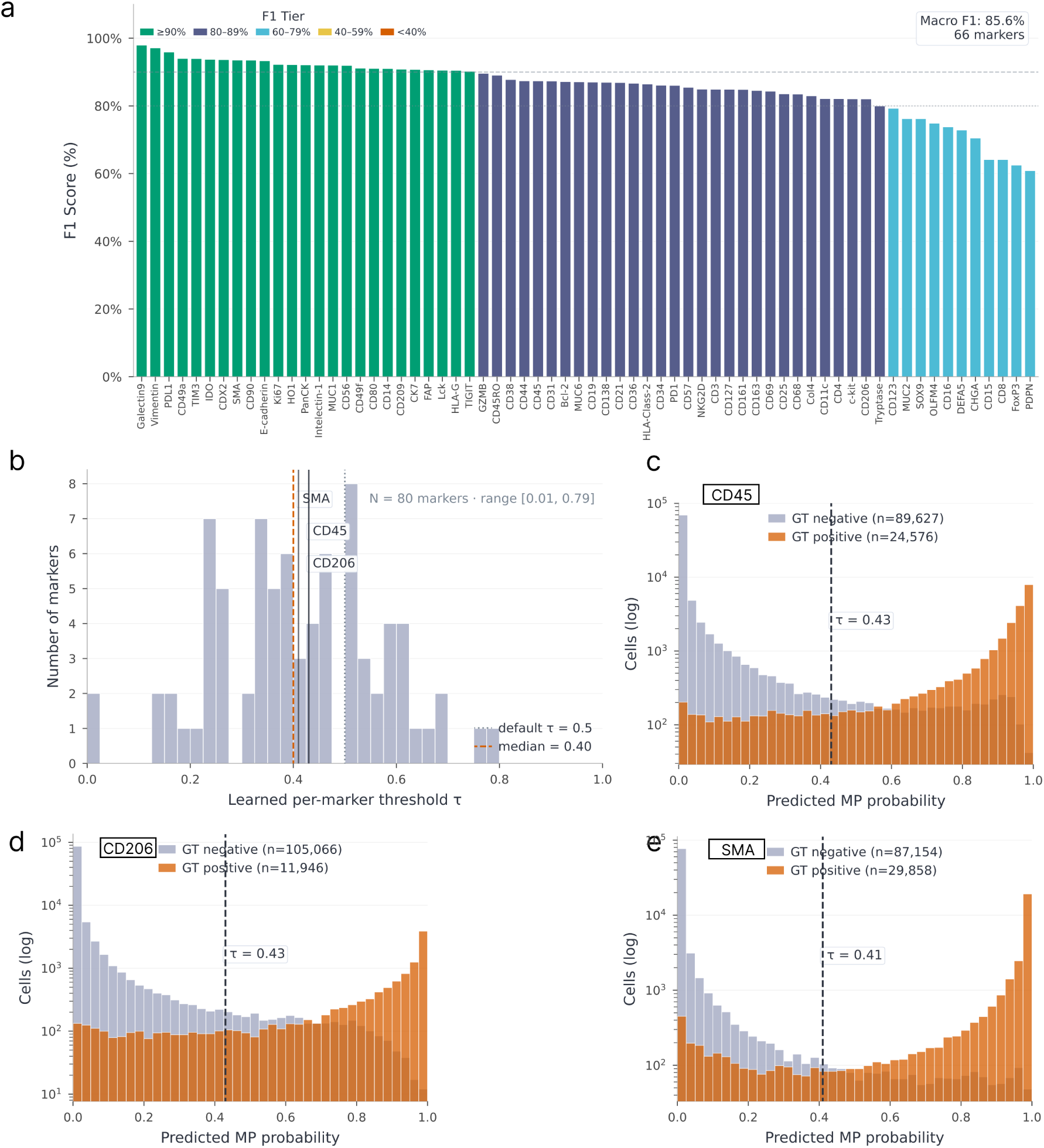
Marker positivity — FiLM head deep dive. **(a)** Per-marker F1 waterfall for DeepCell Types on the held-out test split. Bars are sorted by F1 descending and color-tiered (≥90 / 80–89 / 60–79 / 40–59 / *<*40%). **(b)** Histogram of learned per-marker thresholds *τ* swept on the train split. Most *τ* cluster around 0.3–0.4 (median ≈ 0.40), below the default 0.5, with a long tail of markers calibrating well above or below, justifying the per-marker rather than global threshold. **(c)**–**(e)** FiLM MP decision curves on the test split for CD45 (pan-immune), CD206 (macrophage subset), and SMA (smooth muscle): predicted MP probability stratified by ground-truth positivity, with the learned per-marker *τ* marked. The separation between GT-positive and GT-negative distributions visualizes how marker-specific the head is at its operating threshold.

**Supplementary Figure S8:**
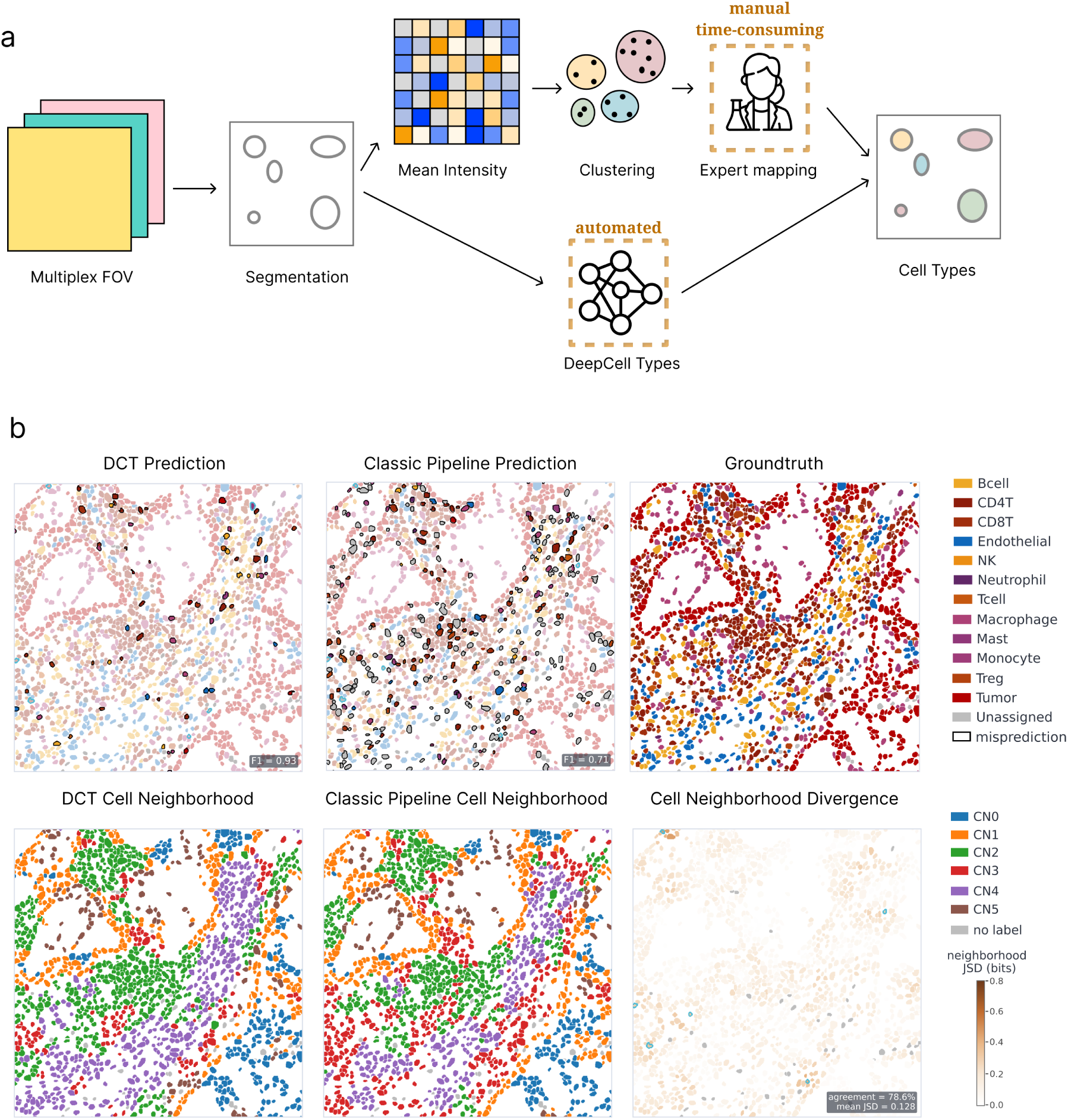
Cell phenotyping workflow comparison. **(a)** Schematic of the two cell-phenotyping workflows. The classic pipeline (per-cell mean intensity → overclustering → expert rule-based mapping) requires a per-dataset gating effort by a domain expert and is time-consuming, whereas the DeepCell Types pipeline is fully automated end-to-end and produces a single set of predictions from one model across all datasets without per-panel adjustments. **(b)** Example FOV evaluated under both pipelines (Sorin LUAD D119 IMC), left to right: DeepCell Types end-to-end predictions, the mean-intensity overcluster (*k* = 200) → expert rule-based merge predictions, and the ground-truth cell types. **(c)** Cellular-neighborhood maps (Schürch et al. 2020; joint KMeans, *k* = 15 nearest-neighbor cell-type composition, shared color key) for the DeepCell Types and traditional labels, and the per-cell Jensen–Shannon divergence between the two workflows’ *k* = 15 neighborhood label distributions (base-2, so values lie in [0, 1] bits). Over the scored cells, the two workflows reach **78.6%** cell-wise label agreement and a low mean JSD of **0.128** bits, indicating that the two predictions are broadly congruent in local neighborhood composition. The expert-driven mapping step in workflow (A) required on the order of hours of hands-on expert time, whereas workflow (B) is fully automated.

**Supplementary Figure S9:**
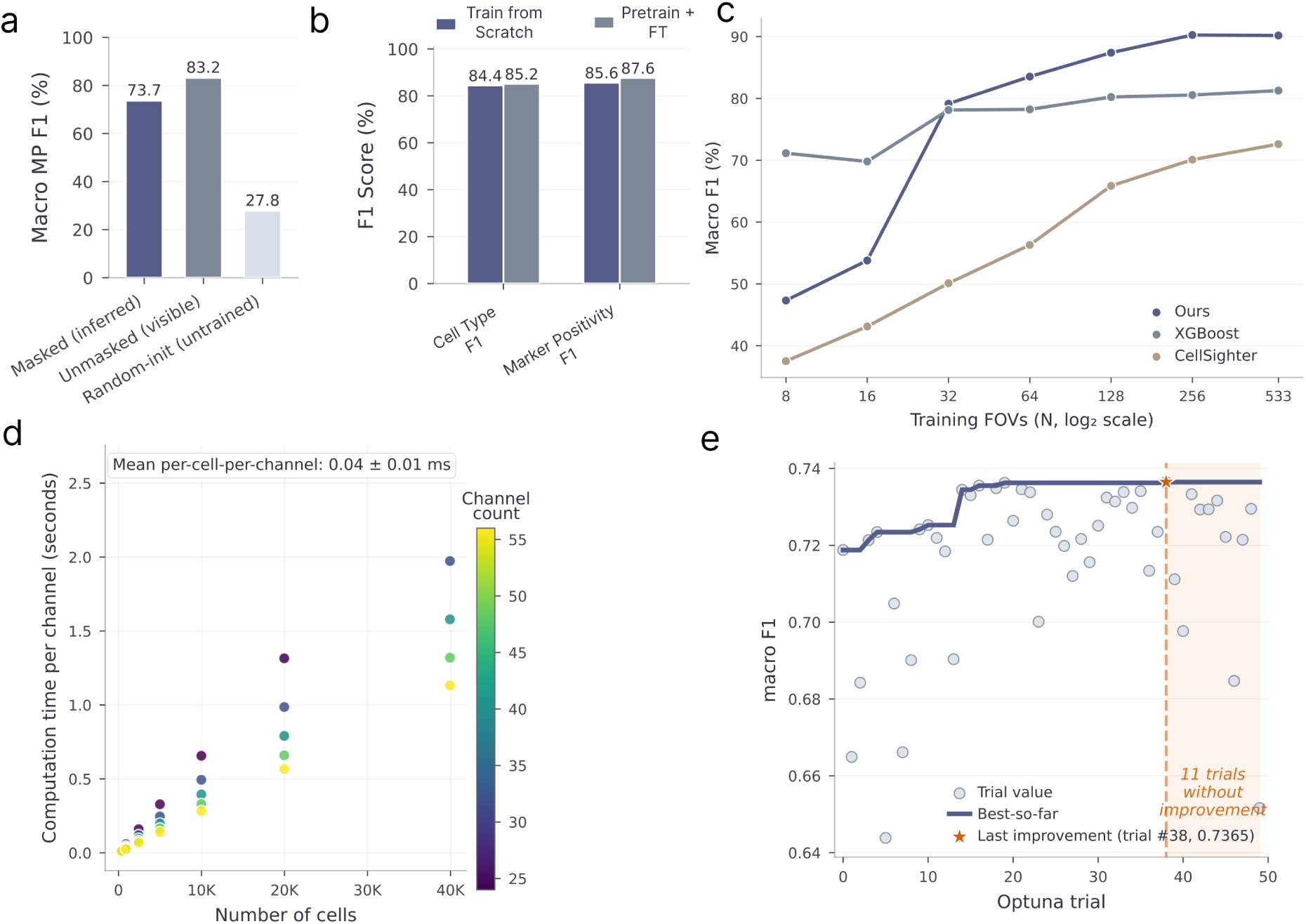
Masked-marker pre-training, annotation efficiency, inference scaling, and baseline tuning. **(a)** Masked-marker pre-training pretext-task diagnostic, complementary to the downstream ablation in panel (b). Macro marker-positivity F1 of the pre-training backbone’s FiLM MP head on the held-out test split under three conditions: *Masked (inferred)*, each marker channel hidden at the input and predicted from the remaining markers and their language embeddings (73.7%); *Unmasked (visible)*, the same head with the marker image visible (83.2%, an upper reference); and *Random-init (untrained)*, the masked setting on an untrained backbone (27.8%, a floor). Masked ≫ random-init and masked ≈ unmasked indicates that the masked objective captures genuine cross-marker structure. **(b)** Self-supervised pretraining ablation. Test-split cell-type macro F1 and marker-positivity F1 for the supervised-from-scratch model versus the masked-marker Pretrain + Fine-tune recipe. Pretraining only provides modest gains on both metrics. **(c)** Annotation-efficiency curve on the Keren MIBI triple-negative breast cancer dataset (*33*): macro F1 as a function of the number of labeled training FOVs (*N*, log_2_ axis), comparing our masked-marker-pretrained backbone (panel b) fine-tuned end-to-end against XGBoost and CellSighter trained from scratch on the same labeled subsets. **(d)** Per-channel inference time scales linearly with the number of cells; on average, computation time per cell per channel is 0.04 ± 0.01 milliseconds. **(e)** Optuna trial progression for the XGBoost baseline hyperparameter sweep on the training partition: each point is one trial’s validation score, with the running best overlaid, confirming the tuned XGBoost baseline reported in Fig. 3c is converged.

### Preprocessing adaptation and fine-tuning for inference on new datasets

Like any supervised model, DeepCell Types is most accurate when inference-time data resemble its training distribution, and the differences that push a new dataset out of distribution — physical resolution, per-channel intensity scaling, and marker naming — are mostly controllable at the preprocessing stage. The released predict interface therefore exposes a preprocess hook that replaces the built-in per-channel normalization without touching model weights, so aligning a dataset is cheap and reversible where retraining is not.

A hand-written preprocess function should respect four requirements. *Resolution*: resample to the training resolution of 0.5 microns per pixel (bilinear interpolation for marker intensities, nearest-neighbor without anti-aliasing for integer segmentation masks). *Intensity* : the built-in step clips each channel at the 99.9th percentile of its non-zero values and min-max scales it within the FOV, so hot pixels that would otherwise set that percentile must be cleaned first. *Marker naming* : harmonize aliases (“Pan-Keratin”/“PanCK”, “CD8”/“CD8a”) to the language-encoder vocabulary — markers outside it are embedded from the templated prompt (table S1) with no retraining — and include canonical lineage markers (CD45, CD31/VWF, CD3/CD4/CD8) for the lineages of interest. *Segmentation*: supply whole-cell masks.

We release this procedure as an installable agent skill (preproc-adapt), automated with the preprocessing adaptor (*54*). It runs a short per-FOV closed loop: pre-register a biological expectation for the tissue and freeze it as a plausibility criterion; run a baseline prediction with a per-channel quality scan; and judge the resulting composition at both the lineage and cell-type level, since a spurious type can hide inside an otherwise-correct lineage. When the composition is implausible, the loop applies one bounded operation from a small declarative set — percentile clipping, log1p, background subtraction, gamma correction, spatial denoising or hot-pixel removal, channel down-weighting or dropping, and a terminal min-max normalization — and re-evaluates, for at most ten rounds. Edits that only inflate confidence or collapse lineage diversity are rejected; an edit is trusted most when it moves multiple lineages toward biology at once, or removes the same confound across tissues with opposite expected compositions — the signature of genuine artifact removal rather than target-steering.

The loop also flags failures preprocessing cannot fix — a model-prior floor, a residual unmoved by principled edits, a coverage gap, or an out-of-distribution modality or panel, alongside classic signals such as a flat per-cell probability distribution or collapse onto a single lineage. These are the cue to label a few representative FOVs in DeepCell Label and fine-tune: cheaply by retraining only the cell-type head on frozen [CLS] embeddings, or — once enough labels are available — end-to-end through the backbone for a higher accuracy ceiling (fig. S9c).

